# Dissociable Cerebellar-Prefrontal Networks Underlying Executive Function: Evidence from the Human Connectome Project

**DOI:** 10.1101/431593

**Authors:** Joseph M. Orr, Trevor B. Jackson, Michael J. Imburgio, Jessica A. Bernard

## Abstract

To date, investigations of executive function (EF) have focused on the prefrontal cortex (PFC), and prominent theories of EF are framed with respect to this brain region. Multiple theories describe a hierarchical functional organization for the lateral PFC. However, recent evidence has indicated that the cerebellum (CB) also plays a role in EF. Posterior CB regions (Crus I & II) show structural and functional connections with the PFC, and CB networks are associated with individual differences in EF in healthy adults. However, it is unclear whether the cerebellum shows a similar functional gradient as does the PFC. Here, we investigated high-resolution resting-state data from 225 participants in the Human Connectome Project. We compared resting-state connectivity from posterior cerebellar ROIs, and examined functional data from several tasks that activate the lateral PFC. Demonstrating preliminary evidence for parallel PFC and CB gradients, Crus I was functionally connected with rostrolateral PFC, Crus II with middle and ventral PFC, and Lobule VI with posterior PFC. Contrary to previous work, the activation of the task thought to activate rostrolateral PFC resembled the connectivity maps of Crus II, not Crus I; similarly, the activation of the task thought to activate middle PFC resembled the connectivity maps of Crus I, not Crus II. Nevertheless, there was evidence for dissociable CB-PFC networks. Further work is necessary to understand the functional role of these networks.

## Introduction

As we navigate our day to day lives, intact executive function (EF) plays a critical role in educational, work, and home settings. EF allows for us to fluidly update our goals and the appropriate behavioral responses online. Indeed, across psychiatric disorders, EF deficits are a prominent feature and contribute to declines in quality of life. EF deficits are thought to be at the core of diseases such as schizophrenia [e.g., Kerns et al., 2008], depression [Snyder, 2013], and substance abuse [Volkow et al., 2011]; this underscores the importance of a clear and complete understanding of this critical cognitive domain. We know that the prefrontal cortex (PFC) plays a crucial role in EF [Miller and Cohen, 2001], and the prominent theories of the neural underpinnings of EF are focused on the PFC.

Nevertheless, the PFC does not act alone, and several cortical (e.g., parietal) and subcortical (e.g., thalamus and striatum) regions have strong connections with the PFC. While PFC-thalamic and PFC-striatal loops are well characterized and integrated into theories of EF [Alexander et al., 1986; Masterman and Cummings, 1997], the connections between the PFC and cerebellum [Bernard et al., 2012; Bernard et al., 2014; Bernard et al., 2016; Kelly and Strick, 2003; Salmi et al., 2010] and their role in EF, are less understood. While the suggestion that the cerebellum may play a role in non-motor behavior is not a new idea [Leiner et al., 1986; Leiner et al., 1993; Schmahmann and Sherman, 1998], more recently there has been an increase in evidence to support this idea more directly. Investigations in non-human primates have revealed distinct closed-loop circuits connecting anatomically segregated regions of the cerebellum and motor and prefrontal cortices, respectively [Dum and Strick, 2003; Kelly and Strick, 2003], and these segregated circuits have also been replicated in the human brain with both structural [Bernard et al., 2016; Salmi et al., 2010] and functional [Bernard et al., 2012; Bernard et al., 2014; Krienen and Buckner, 2009; O’Reilly, 2010] measures of brain connectivity. While some investigations have directly compared connectivity between lobules at rest, these investigations have focused on sensorimotor function [Kipping et al., 2013] or associations with known cortical networks [Sang et al., 2012]. Importantly however, Sang and colleagues did suggest heterogeneity in the connectivity patterns of the posterior aspects of the cerebellum. Further, meta-analytic connectivity modelling has suggested co-activation between Crus I and II of the cerebellum and prefrontal cortical regions during task performance [Balsters et al., 2014] providing further support for the purported importance of these regions in EF. Direct comparisons of the connectivity patterns of Crus I and II are thus warranted for our understanding of the cerebellum in EF, and to better elucidate whether a similar functional processing gradient exists in both the cerebellum and PFC.

Paralleling the closed-loop circuitry of the cerebellum, lateral posterior regions of the cerebellum have been implicated in cognitive processing. Early work in patients with cerebellar lesions demonstrated both cognitive and affective deficits in patients with posterior cerebellar lesions [Schmahmann and Sherman, 1998]. More recently, work using functional imaging has suggested a corresponding functional topography in the cerebellum [e.g., E et al., 2012; Stoodley et al., 2012; Stoodley and Schmahmann, 2009]. Lateral posterior lobules of the cerebellum (Crus I, Crus II), show activation during the performance of cognitive tasks, including EF tasks. This parallels the regions of the structure showing both structural and functional connections with the PFC [Bernard et al., 2012; Bernard et al., 2016; Krienen and Buckner, 2009; O’Reilly et al., 2010; Salmi et al., 2010]. However, the specific contribution these posterior lobules make to cognitive behavior remains unknown. It has been suggested that much like in the motor domain, Crus I and Crus II of the cerebellum process internal models of thought, that aid in, and allow for organized and efficient cognition [Ito, 2008; Ramnani, 2006]. More recently, Ramnani [2014] has suggested that connections between the PFC and cerebellum support the trade-off or transition during cognitive learning from top-down control to automatic processing according to internal models. With that said, it may be the case that these regions differentially contribute to distinct components of EF, subserved by their connectivity patterns with the prefrontal cortex.

There are a number of theories describing the functional organization of the PFC. Many of these theories suggest that the PFC is organized in a rostral-caudal gradient of abstraction [Badre, 2008; O’Reilly, 2010], and as such, different regions of the PFC contribute to different components of EF. The most anterior aspects of the PFC (rostral lateral prefrontal cortex; RLPFC) are thought to be involved in abstract higher order behaviors like the control of multiple goals, prospective memory, or using internal goals to guide task selection [Burgess et al., 2007; Koechlin and Hyafil, 2007; Orr and Banich, 2014]. Conversely, the more posterior regions of the PFC (i.e., premotor cortex and inferior frontal gyrus) are thought to be involved in more concrete behaviors related to action, including sensory or response selection [Banich, 2009]. While some accounts suggest this abstraction gradient is hierarchical (with the RLPFC being at the apex), more recent evidence suggests that the dorsolateral PFC (DLPFC) is the apical region of the PFC. Regardless of the nature of this organization, intact PFC function is thought to be necessary for maintaining healthy EF. However, the PFC is unlikely to be acting alone. A network-based approach to understanding EF is crucial, and in particular, an eye towards the cerebellum is warranted.

To investigate cerebellar-prefrontal networks, we utilized data from 225 neurotypical unrelated participants in the Human Connectome Project (HCP), due to its large size and exceptional data quality—ultra-high resolution (0.7 mm isotropic) structural images, high-resolution fMRI scans (3 mm isotropic)—from a variety of cognitive, emotion, and motor tasks, and multiple sessions of resting-state fMRI (rfMRI). Data in the HCP comes from a healthy young adult (21-35 years) community sample. We focused on rfMRI data for the current study in order to identify dissociable cerebellar-prefrontal networks. Cerebellar regions-of-interest were defined from the SUIT Probabilistic atlas [Diedrichsen et al., 2009]. We selected three lateral posterior cerebellar regions from the left and right hemispheres: Crus I, Crus II, and Lobule VI. Lobule VI was chosen as it has been shown to be functionally connected to posterior prefrontal cortex [Bernard et al., 2012]. We predicted that regions of the lateral posterior cerebellum would show connectivity patterns with the prefrontal cortex that parallel the rostral-caudal gradient of abstraction, suggesting that these lobules may subserve more specific components of EF.

In order to gain additional insight into the nature of CB-PFC network contributions to EF, we also examined task-based fMRI. Tasks in the HCP targeted a variety of functional domains including motor, language, working memory, relational processing, social processing, emotional processing, and gambling. These tasks were chosen to activate as many different functional network nodes as possible, in order to serve as seeds for connectivity analyses [Barch et al., 2013]. We chose four tasks that are reliable localizers for lateral prefrontal regions: the Relational Processing Task, the N-Back Working Memory Task, a Language Comprehension Task, and Motor Tapping.

The Relational Processing Task has been proposed as a localizer task for the RLPFC [Smith et al., 2007]. In this task, participants compared items on simple features (control condition) or determined if the same feature changed between two sets (i.e., relational condition). We predicted that the Relational Processing Task would activate the RLPFC and Crus I networks to a greater degree than the other tasks. The N-Back Task was embedded in a category representation task that has been shown to be a reliable localizer for the DLPFC [Barch et al., 2013; Drobyshevsky et al., 2006]. Therefore, we predicted that the Working Memory Task would activate the DLPFC and Crus II networks to a greater degree than the other tasks. The Language Comprehension Task involved comparing auditory stories with comprehension questions to math problems [Binder et al., 2011]. This task has been shown to activate inferior frontal and temporal regions associated with language processing. In the cerebellum, language tasks have been shown to activate both Crus I and Lobule VI [Stoodley et al., 2012; Stoodley and Schmahmann, 2009]. The Motor Mapping Task involved simple motor movements of the hands, toes, and tongue. We chose the right hand condition, which should activate the left premotor and motor cortices, and right cerebellar lobules IV-V and VIII [Stoodley et al., 2012].

A recent paper analyzed functional task activation in the cerebellum in these same tasks (using a less restricted set of HCP data), and found evidence for three distinct nonmotor representations [Guell et al., 2018a]. They found that working memory was associated with Crus I, Crus II, and Lobule VIIIb, language was associated with activation in Crus I and Crus II (more medial than the working memory activation), and right hand tapping activation was associated with Right Lobules V-VI and VIII. In their main manuscript they state that the relational processing task did not yield sufficient activation, but in their supplementary materials they report a sub-threshold activation map for the relational processing task; this revealed that the relation vs. control match comparison was associated with similar activation as the working memory task.

Critically, Guell and colleagues [2018a] took a different approach than the current work. First, they only examined cerebellar task activation. Our goal was to identify dissociable cerebellar-PFC networks, while the goal of Guell and colleagues was to identify a functional topography in the cerebellar. Second, Guell and colleagues included participants irrespective of education, drug use history, and personal and family mental health history, while we selected “clean” controls with high levels of education, no history of drug use, and no personal or family mental health history. Third, we only chose unrelated individuals, as including related individuals violates statistical assumptions of independence [Winkler et al., 2014]. Lastly, Guell and colleagues [2018a] did not perform a group-level GLM, rather they calculated Cohen’s d based on the average contrast maps, and chose a threshold of Cohen’s d > 0.5. We performed a statistically conservative GLM using permutation methods, and for task-based activation, used grayordinate data (representing cortical data as a surface and subcortical/cerebellar data as a volume) which shows better cortical localization [Coalson et al., 2018].

## Methods

### Participants – HCP

Structural, resting-state fMRI (rfMRI), and cognitive task fMRI (tfMRI) data from the S900 Release of Human Connectome Project (WU-UMN HCP Consortium) were used in this investigation. The goal of recruitment in the WU-UMN HCP Consortium was to capture a broad range of variability in healthy individuals with respect to behavioral, ethnic, and socioeconomic diversity [Van Essen et al., 2012]. Lifestyle and demographic data were collected alongside the imaging data and were used in this study to select for a sample meant to be representative of unrelated, non-clinical individuals across a variety of socioeconomic, behavioral, and ethnic backgrounds in order to maintain generalizability and control for any potential structural and functional similarities and differences linked to the factors above. Detailed descriptions of each variable used to eliminate participants are available here: https://wiki.humanconnectome.org (see HCP Data Dictionary Public- Updated for the 1200 Subject Release). To establish an ideal picture of CB-PFC network functioning without the influence of potential drug or disease confounds, we only selected data from unrelated individuals who psychologically and neurologically “clean”: right-handed, high-school graduate or greater, no family history of mental illness, and no alcohol or drug abuse. More specifically, data were considered for this study only if the participant displayed right-handedness (Handedness > 24), attained a high school degree (SSAGA_Educ > 11), reported no family history of mental illness (FamHist_*_None = 1), did not meet the DSM4 criteria for Alcohol Abuse or Dependence (SSAGA_Alc_D4_Ab_Dx ! = 5; SSAGA_Alc_D4_Dp_Dx ! = 5), and did not meet the DSM criteria for Marijuana Dependence (SSAGA_Mj_Ab_Dep = 0). Data was further excluded if the participant reported more than 7 drinks per week for a female or 14 drinks per week for a male ([F]Total_Drinks_7days <8 OR [M]Total_Drinks_7days <15). Only one randomly selected participant from each family unit was used in order to account for any potential similarities in brain structure and function [Ganjgahi et al., 2015; Glahn et al., 2007; Pagliaccio et al., 2015; Thompson et al., 2001]. These exclusions resulted in a sample size of 225 individuals ranging in age from 22 to 36 years (92 males, 133 females; see Supplemental Material for a list of participants included).

### HCP image acquisition and preprocessing

Details on data acquisition in the HCP sample are reported by Van Essen and colleagues [2012]. rfMRI data for each participant consisted of 2, 15-minute scans (1200 volumes, 720 ms TR, 2 mm isotropic voxels); although the HCP database includes up to 4 rfMRI scans per each participant with 2 pairs of scans collected on different days, we selected only one pair of scans primarily to restrict demands on data storage, but also to avoid any potential fluctuations in cognitive state across the different days [Poldrack et al., 2015]. rfMRI data came from the Resting State fMRI 1 FIX-Denoised (Extended) Package which included preprocessed volumetric timeseries data that had been denoised using the FIX ICA-based automated method and registered with Multimodal Surface Matching-Folding (MSMsulc), a technique that results in superior registration of functional and structural data using cortical folding [Robinson et al., 2014]. Additional details on this pipeline are discussed in detail elsewhere [Glasser et al., 2013]. Note, that the data resulting from the HCP preprocessing was not smoothed, and any smoothing described below was on denoised, but unsmoothed data.

tfMRI data for each participant consisted of cross-run analyzed grayordinates data, a format that represents gray matter locations by cortical surface vertices or subcortical volume voxels. tfMRI data had undergone preprocessing with the *fMRISurface* pipeline [Glasser et al., 2013], registration using the advanced MSMall method (which utilizes cortical folding and function to align data), and were then analyzed via GLM through the first-level (individual run level) and second-level (across runs, within subject) with the HCP Task fMRI Analysis Pipeline [Barch et al., 2013]. The grayordinates-based analysis is similar to a traditional volume-based analysis, with a GLM implemented in FSL using autocorrelation correction [Woolrich et al., 2001], however, spatial smoothing was done on the left and right hemisphere surfaces using a geodesic Gaussian algorithm. Subcortical gray matter time series were smoothed within defined gray matter parcels. Smoothing by 2 mm FWHM was done by the HCP *fMRISurface* Pipeline, and an additional smoothing was done in subsequent analyses to bring the total smoothing to 4 mm FWHM.

In addition to functional data, preprocessed T1 structural data were also downloaded for each participant. Structural scans had undergone gradient distortion correction, bias field correction, and registration to the 0.8 mm resolution MNI brain using MSMsulc. Structural images were used for tissue type segmentation for purposes of rfMRI data processing (see below).

### Data analysis

#### Resting-state Functional Connectivity Analysis

rfMRI data underwent all additional processing and analysis using the CONN toolbox [v. 17e; Whitfield-Gabrieli and Nieto Castañón, 2012], a Matlab-based application designed for functional connectivity analysis. CONN was compiled as a standalone application for MATALB R2016b in centOS 6 running on a 128-core Intel Xeon Broadwell blade cluster. Preprocessing in CONN consisted of structural segmentation, smoothing (6 mm FWHM), and artifact detection (global signal z-value threshold: 5, subject motion threshold: 0.9 mm). Data were then denoised with linear regression with confound regressors for 5 temporal components each from the segmented CSF and white matter, 24 motion realignment parameters, signal and/or motion outliers, a linear trend regressor, and the 1st order derivative from the effect of rest. Although the data had already been denoised using ICA, we took an aggressive approach towards denoising and included the motion parameters (determined before ICA) in this regression. Finally, data underwent bandpass filtering (0.01–0.1 Hz).

Functional connectivity from Left and Right Crus I, Crus II, and Lobule VI to the rest of the brain was examined using a General Linear Model. ROIs were defined from the SUIT probabilistic cerebellum atlas [Diedrichsen et al., 2009]. The location of these ROIs are shown in Figure 8. The average BOLD signal timeseries was extracted from each ROI. At the first-level, a standard GLM was performed where the canonical hemodynamic response function was convolved with the rest condition. A semi-partial correlation approach was used, where all 6 ROIs were entered into one model to estimate their unique contributions. At the group-level, βs were saved as Fisher-transformed correlation coefficients. The connectivity of each ROI was considered against the other two ROIs in pairwise contrasts, e.g., Crus I > Lobule VI was coded as [1, 0, −1]. Thus, 6 contrasts were defined: Crus I > Crus II, Crus I > Lobule VI, Crus II > Crus I, Crus II > Lobule VI, Lobule VI > Crus I, Lobule VI > Crus II. Statistical maps were thresholded non-parametrically using a conservative thresholding approach, given the very large sample size; the cluster defining threshold was set to an FDR-corrected p < .001, and the resulting clusters were thresholded to a cluster-mass FDR-corrected p < .001 with 10,000 permutations. To better illustrate the patterns of connectivity for each ROI relative to the other two, we generated conjunction maps for each ROI e.g., (Crus I > Crus II) Λ (Crus I > Lobule VI). This was accomplished by creating binary masks of the corrected contrast maps and multiplying the two appropriate masks together. Cluster tables were generated using FSL’s autoaq, a tool which performs automatic atlas queries for the center of mass for each cluster. We used the Harvard-Oxford Cortical and Subcortical Atlases, and for prefrontal clusters, we used the Sallet Dorsal Frontal Atlas [Sallet et al., 2013] and the Neubert Ventral Frontal Atlas [Neubert et al., 2014]. As these atlases were only available for the right hemisphere, to label left hemisphere clusters we flipped the statistical maps along the x-axis. The volumetric thresholded results were mapped to the cortical surface to allow for better comparison with the task fMRI analyses. The Connectome Workbench (v. 1.3.1; https://www.humanconnectome.org/software/connectome-workbench) was used to map the volumes to the HCP S1200 Group Average Inflated Surface [Van Essen et al., 2017].

#### Task fMRI Group Analysis

As mentioned above, tfMRI data were downloaded in a pre-analyzed format, which consisted of cross-run analyzed dense grayordinates data smoothed to 4 mm. Contrast Parameter Estimate (COPE) images from each participant were selected from each task for inclusion in group-level analyses. For the relational processing task, we chose the contrast for Relational > Match; for the working memory task, we chose the contrast for 2-back > 0-back; for the language task, we chose the contrast for Story > Math; and for the motor task we chose the contrast for Right Hand > Average. Data were analyzed with the Connectome Workbench. Analysis scripts are available at {create repository}. For each task, the COPE maps from each participant were merged and the merged files were then separated into volumetric data for subcortical areas and surface data for the left and right hemispheres, resulting in three maps for each task. For the left and right hemispheres, the midthickness surface mesh was used so that the adjacency (neighborhood) information between vertices can be computed. Group means for each task were then analyzed using non-parametric permutation testing in PALM [Winkler et al., 2014] with 5000 permutations for each test. Clusters were identified with Threshold Free Cluster Enhancement (TFCE) [Smith and Nichols, 2009] with FWE-corrected *p*-values, stored as -*log(p)* values to increase the spread of volumes for viewing purposes. Because 3 tests (left and right hemisphere surface and subcortical volume) were run for each of the four tasks, we set a minimum threshold of -*log(p)* = 2.38 which is equivalent to .004166 or .05/12. As this threshold produced large areas of overlap, and we wanted to identify unique clusters of activation, we further thresholded the maps at the 95^th^ percentile of the t-values. As there are currently no tools available to create cluster tables in Connectome Workbench, we identified the location of activation clusters by overlaying the results on the label file of the Glasser et al. [2016] Multi-Modal Parcellation of 180 cortical areas. We placed a vertex in the center of each cluster of activation, and for large clusters that covered multiple areas, a vertex was placed in each area covered by the activation. The results (including the Multi-Modal Parcellation label file) are available on the Brain Analysis Library of Spatial Maps and Atlases (BALSA; https://balsa.wustl.edu/study/show/k53g) and on the Open Science Framework (https://osf.io/3vka4). Analysis scripts for the task fMRI analyses are also available on the Open Science Framework.

## Results

### Resting-State Functional Connectivity Results

#### Crus I

Connectivity from each ROI to the rest of the brain was contrasted with the other ROIs. Examining the conjunction of the contrasts of Crus I > Crus II and Crus I > Lobule VI, Crus I showed greater connectivity with several prefrontal regions: the RLPFC, the frontal operculum, the rostral cingulate, and the dorsal premotor cortex (see Figure 1 and Tables 1&2). There were some notable differences between the left and right cerebellar ROIs. While there were bilateral RLPFC clusters for left Crus I, only the left RLPFC showed connectivity with right Crus I. In addition, the frontal operculum connectivity was only present for right Crus I. We further explore these discrepancies below.

**Figure 1.**
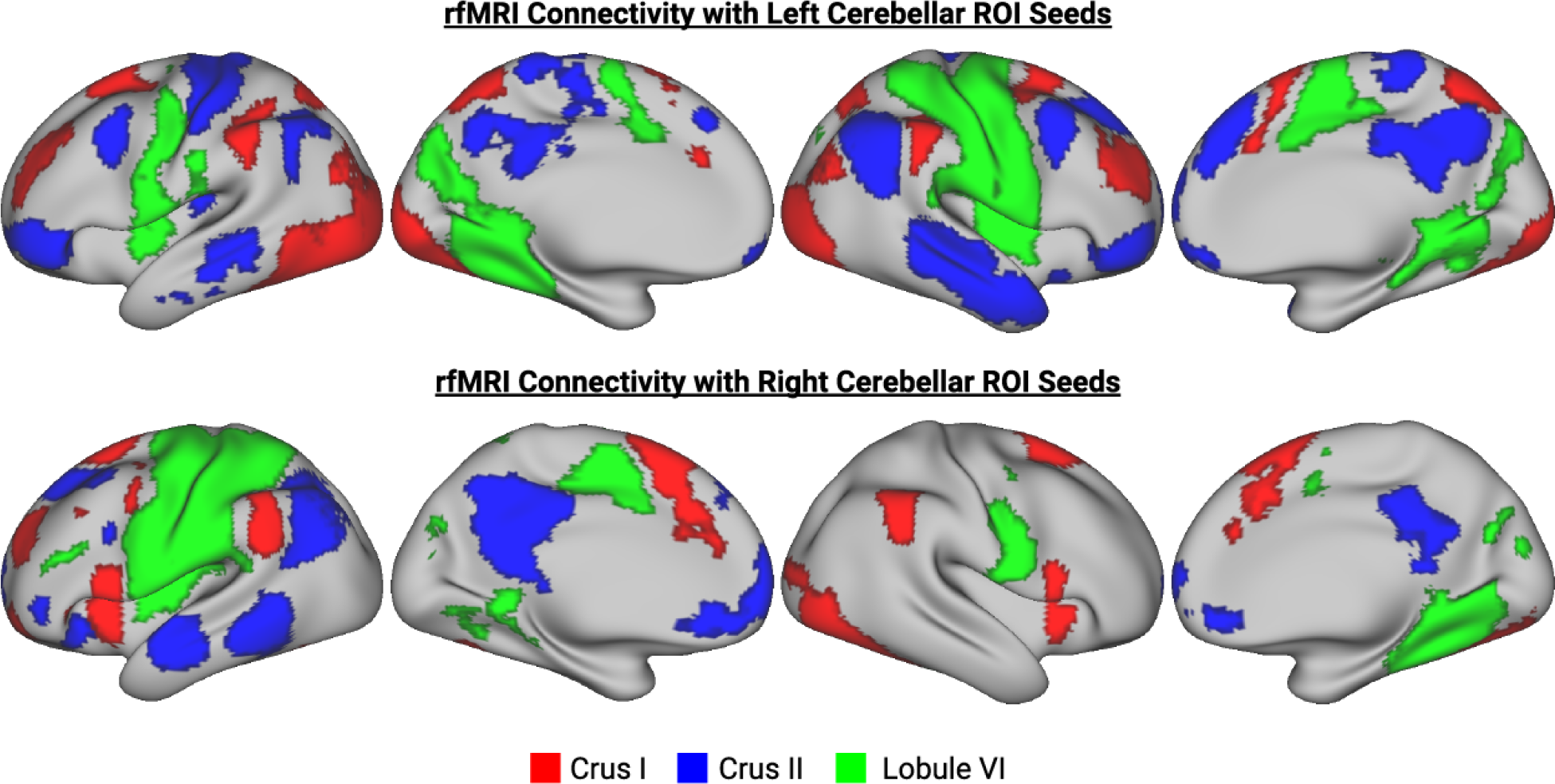
Peak activations from the four functional tasks: working memory (red), relational processing (blue), language (green), and motor (yellow) overlaid on the inflated cortical surface (top row in each section) and the cerebellum (bottom row in each section). Cluster-corrected activation maps were further thresholded at the 95 percentile to isolate peaks of activation, and were binarized for display. The working memory activation was thresholded at T > 10.98, relational processing was thresholded at T > 9.34, language was thresholded at T > 15.5, and motor was thresholded at T > 13.45.

**Table 1.** Atlas query of conjunction of Left Crus I connectivity results. For large clusters, sub-region labels are listed below. COG: Center of Gravity.

|  |  |  |  |  |  |
| --- | --- | --- | --- | --- | --- |
| Cluster Index | Voxels | COG X (mm) | COG Y (mm) | COG Z (mm) | COG Label |
| 10 | 10476 | -17.4 | -79.5 | -13.9 | 42% Occipital Fusiform Gyrus |
| 9 | 2175 | 10.2 | -68.8 | 51.5 | 42% Precuneous Cortex |
| 8 | 865 | 15.2 | 15.3 | 59.6 | 47% PreSMA |
| 7 | 619 | -20.5 | 10.1 | 62.4 | 45% Ant PMd |
| 6 | 596 | 41.6 | -47.5 | -44.2 | 73% Right Crus II |
| 5 | 544 | -49.7 | -42.5 | 46.4 | 40% Supramarginal Gyrus |
| 4 | 538 | 31.6 | 50.9 | 29 | 51% Area 46 |
| 3 | 520 | -28.4 | 48.2 | 29 | 61% Area 9/46D |
| 2 | 495 | 51.7 | -39.6 | 44.1 | 52% Supramarginal Gyrus |
| 1 | 27 | 12.5 | -102 | 16.7 | 51% Occipital Pole |
| ----- |  |  |  |  |  |
| Structures to which each cluster belongs to: |  |  |  |  |  |
| Cluster #10 |  | Average Atlas Probability Within Cluster |  |  |  |
| Lateral Occipital Cortex, inferior division |  | 10.4 |  |  |  |
| Occipital Fusiform Gyrus |  | 4.9 |  |  |  |
| Occipital Pole |  | 15.2 |  |  |  |
| Cluster #9 |  | Average Atlas Probability Within Cluster |  |  |  |
| Lateral Occipital Cortex, superior division |  | 40.8 |  |  |  |
| Precuneous Cortex |  | 11.3 |  |  |  |
| Cluster #8 |  | Average Atlas Probability Within Cluster |  |  |  |
| Supplementary Motor Area |  | 9.9 |  |  |  |
| Pre-Supplementary Motor Area |  | 8.5 |  |  |  |
| Area 8A |  | 10.6 |  |  |  |
| Anterior Premotor Dorsal (PMd) |  | 13.6 |  |  |  |
| Cluster #7 |  | Average Atlas Probability Within Cluster |  |  |  |
| Pre-Supplementary Motor Area |  | 7.0 |  |  |  |
| Anterior Premotor Dorsal (PMd) |  | 29.6 |  |  |  |
| Cluster #6 |  | Average Atlas Probability Within Cluster |  |  |  |
| Right Crus I |  | 22.0 |  |  |  |
| Right Crus II |  | 22.3 |  |  |  |
| Right VIIb |  | 14.1 |  |  |  |
| Cluster #5 |  | Average Atlas Probability Within Cluster |  |  |  |
| Supramarginal Gyrus, anterior division |  | 19.8 |  |  |  |
| Supramarginal Gyrus, posterior division |  | 28.1 |  |  |  |
| Cluster #4 |  | Average Atlas Probability Within Cluster |  |  |  |
| Area 9/46D |  | 20.9 |  |  |  |
| Area 46 |  | 18.0 |  |  |  |
| Cluster #3 |  | Average Atlas Probability Within Cluster |  |  |  |
| Area 9/46D |  | 29.0 |  |  |  |
| Area 46 |  | 26.6 |  |  |  |
| Area 8B |  | 7.9 |  |  |  |

**Table 2.**
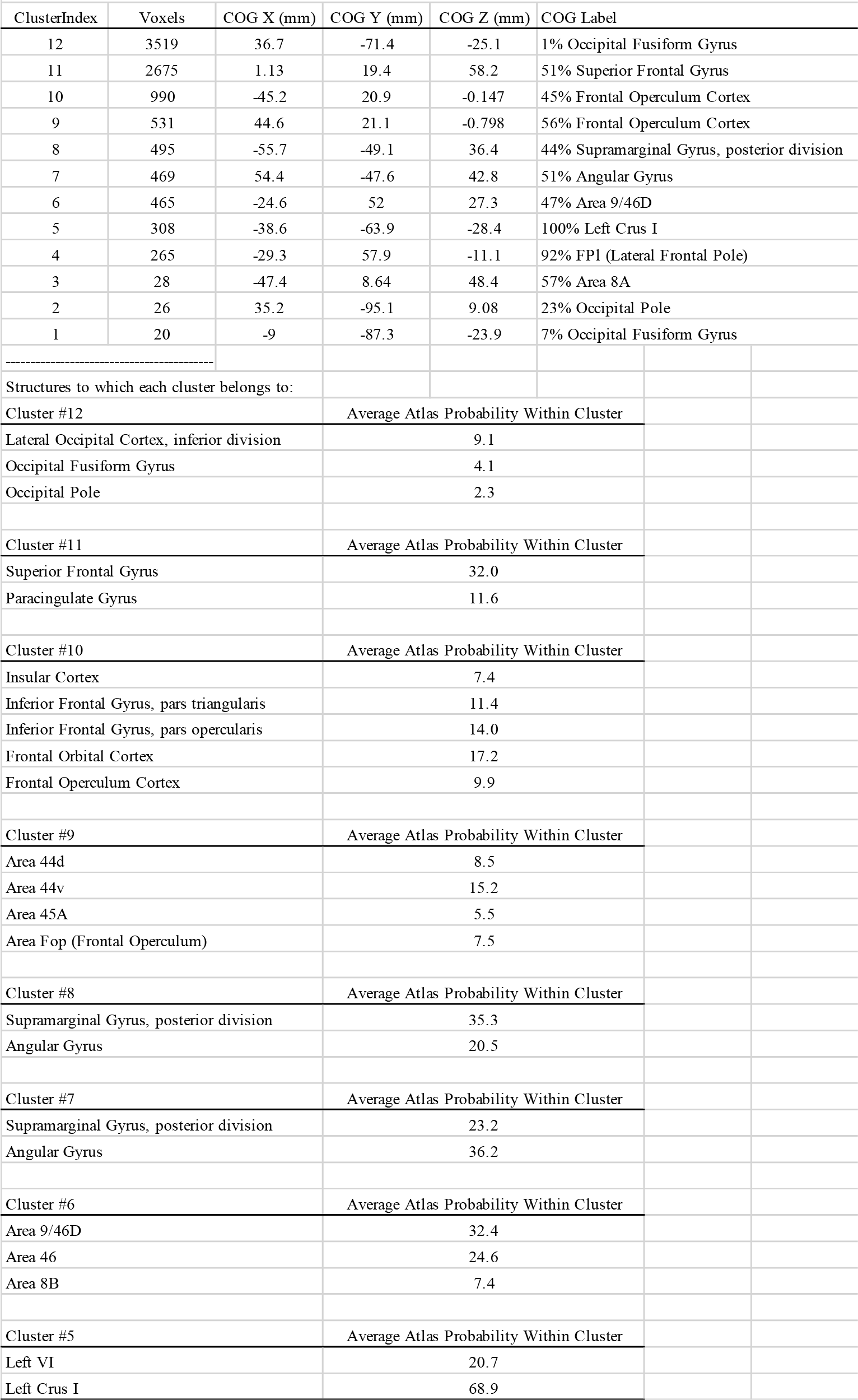
Atlas query of conjunction of Right Crus I connectivity results. For large clusters, sub-region labels are listed below. COG: Center of Gravity.

In the RLPFC, Crus I connectivity was localized to area 46 and area 9/46d. While area 46 is often referred to as frontopolar area 10 [Gilbert et al., 2010; Leung et al., 2005], work has shown that this region differs from the more anterior area 10 in terms of its cytoarchitectonics [Bludau et al., 2013] as well as structural connectivity and functional coupling [Sallet et al., 2013]. Sallet and colleagues [2013] showed that area 46 shows maximal functional coupling with dorsal premotor cortex, rostral anterior cingulate, and inferior parietal lobule, while area 10 shows maximal coupling with ventromedial PFC and amygdala. As shown in Figures 2 & 3 (see also Supplemental Tables 1-4), the direct comparison of Crus I with each of the other two ROIs supported the results of the conjunction analysis. The main difference between the conjunction analyses and the direct comparison was the finding that the frontal operculum was only present in the comparison of Left Crus I and II, but not in the comparison of Left Crus I and Lobule VI. With the exception of the frontal operculum, the connectivity maps for the direct comparisons of left and right Crus I were more similar than suggested by the conjunction analyses. Overlaying the conjunction connectivity maps on the Yeo 17-area parcellation [Yeo et al., 2011] revealed that the RLPFC cluster, the rostral cingulate cluster, and the frontal operculum cluster were all part of the cingulo-opercular task control network, a network involved in stable task-level control and maintenance of goals [Dosenbach et al., 2008] (See Figure S1). Thus, as predicted, Crus I shows greater connectivity with regions of prefrontal cortex involved in higher-level abstract control compared to Crus II and Lobule VI.

**Figure 2.**
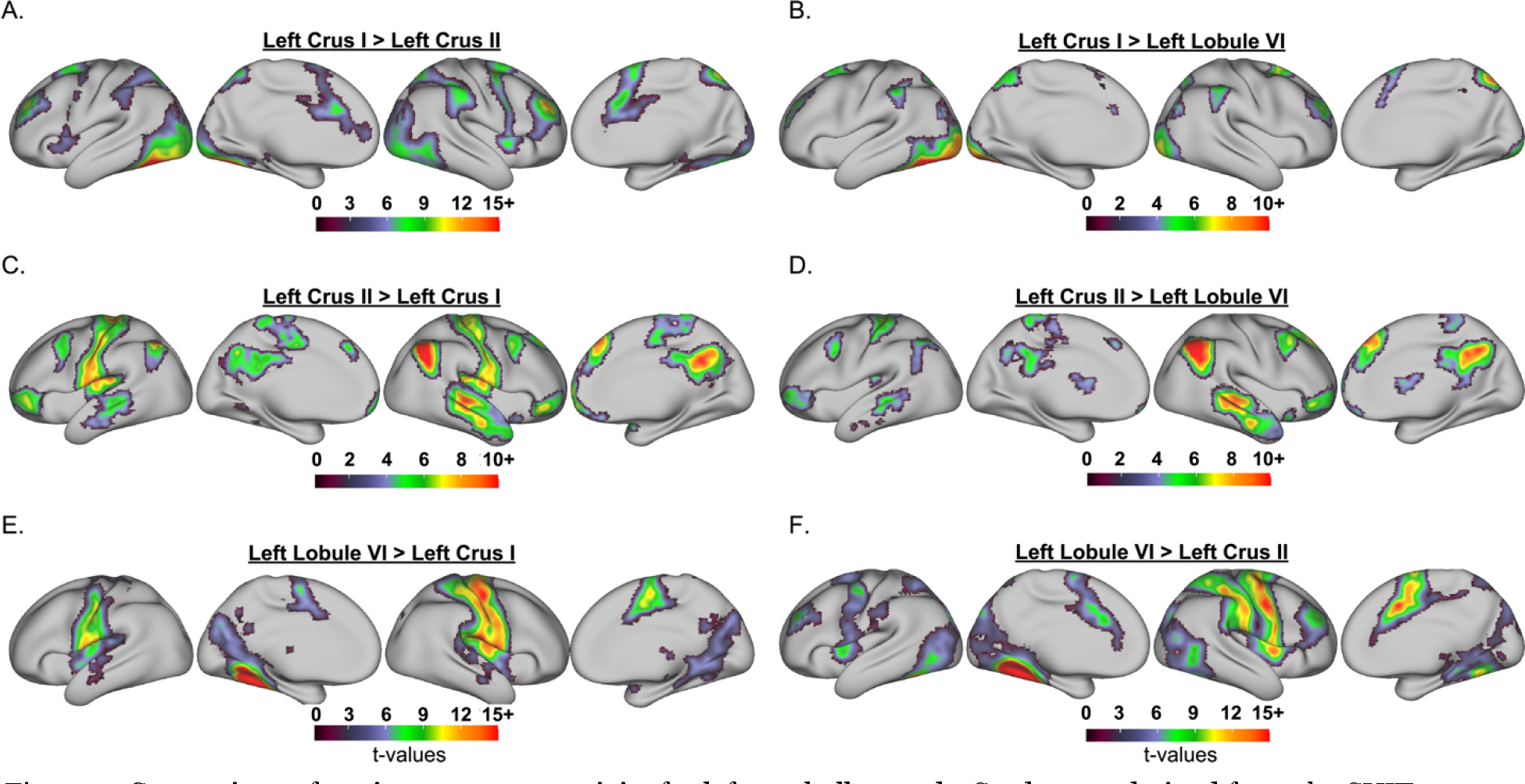
Comparison of resting-state connectivity for left cerebellar seeds. Seeds were derived from the SUIT Probabilistic Cerebellar Atlas and included Crus I, Crus II, and Lobule VI. Pairwise comparisons for seed-to-voxel connectivity were created with linear contrasts in CONN that included all three seeds and setting the weight for the third seed that was not of interest to 0, e.g., for the contrast of Left Crus I > Left Crus II, the weight for Left Lobule VI was set to 0, i.e., [1 −1 0].

**Figure 3.**
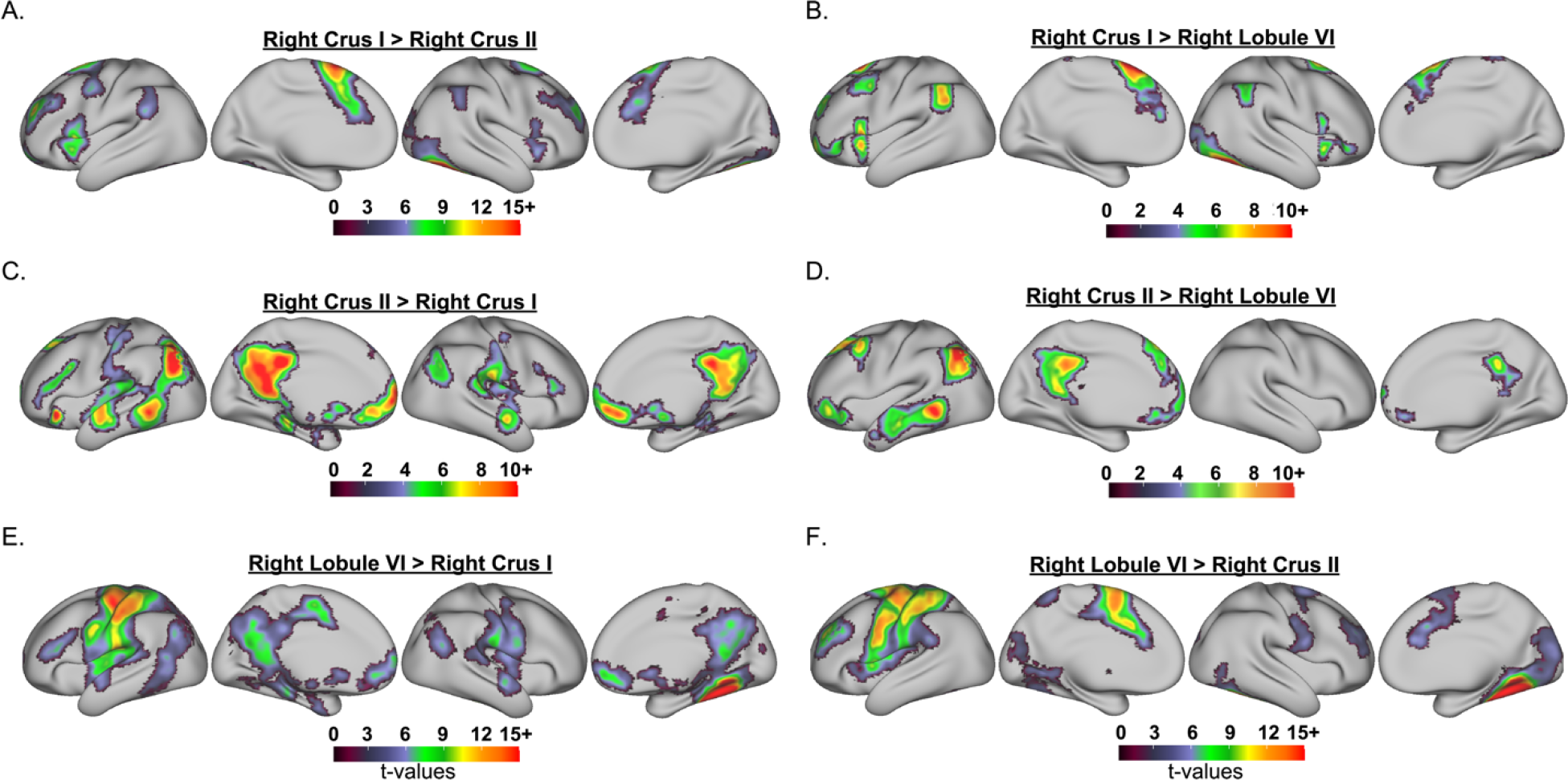
Comparison of resting-state connectivity for right cerebellar seeds. Seeds were derived from the SUIT Probabilistic Cerebellar Atlas and included Crus I, Crus II, and Lobule VI. Pairwise comparisons for seed-to-voxel connectivity were created with linear contrasts in CONN that included all three seeds and setting the weight for the third seed that was not of interest to 0, e.g., for the contrast of Right Crus I > Right Crus II, the weight for Right Lobule VI was set to 0, i.e., [1 −1 0].

#### Crus II

Turning to Crus II, the conjunction of Crus II > Crus I and Crus II > Lobule VI revealed two main areas of connectivity in the prefrontal cortex, one posterior DLPFC cluster, and one anterior ventrolateral PFC cluster (See Figure 1 and Tables 3 & 4). The posterior DLPFC cluster was located in areas 8A and ventral 9/46 in the left hemisphere and areas 8B, 9, and pre-SMA in the right hemisphere. The second area of connectivity was located in the Inferior Frontal Sulcus and area 47. Area 47 has been shown to exhibit functional coupling with nearby regions of anterior prefrontal cortex, precuneus, as well as anterior lateral temporal cortex [Neubert et al., 2014]. Indeed, anterior lateral temporal cortex and precuneus were also connected with Crus II.

**Table 3.** Atlas query of conjunction of Left Crus II connectivity results. For large clusters, sub-region labels are listed below. COG: Center of Gravity.

| ClusterIndex | Voxels | COG X (mm) | COG Y (mm) | COG Z (mm) | COG Label |
| --- | --- | --- | --- | --- | --- |
| 15 | 4622 | -10.1 | -40 | 51 | 35% Postcentral Gyrus |
| 14 | 2876 | 60 | -15.7 | -16.1 | 45% Middle Temporal Gyrus, posterior division |
| 13 | 2781 | -25.8 | -76.8 | -38.4 | 88% Left Crus II |
| 12 | 2397 | 15.9 | 41 | 40.8 | 55% Area 8B |
| 11 | 1591 | 49.4 | -58.4 | 35.7 | 36% Angular Gyrus |
| 10 | 1349 | 28.7 | -79 | -33.8 | 91% Right Crus I |
| 9 | 1211 | -23.6 | 55.1 | -7.96 | 96% FPI (Lateral Frontal Pole) |
| 8 | 720 | 45.1 | 46 | -8.95 | 55% Area 47 |
| 7 | 612 | -63.3 | -33.8 | -7.68 | 76% Middle Temporal Gyrus, posterior division |
| 6 | 563 | -43.1 | -64.6 | 44.7 | 54% Lateral Occipital Cortex, superior division |
| 5 | 313 | -43.1 | 16.7 | 42.1 | 39% Area 9/46V |
| 4 | 281 | -3.38 | -56.4 | -47.1 | 99% Left IX |
| 3 | 174 | -37.5 | -22.8 | 14.9 | 25% Central Opercular Cortex |
| 2 | 45 | -63.6 | -5.64 | -27.1 | 17% Middle Temporal Gyrus, anterior division |
| 1 | 39 | 33.7 | 21.6 | -20.8 | 99% Fop (Frontal Operculum) |
| Structures to which each cluster belongs to: |  |  |  |  |  |
| Cluster #15 |  | Average Atlas Probability Within Cluster |  |  |  |
| Precentral Gyrus |  | 11.6 |  |  |  |
| Postcentral Gyrus |  | 16.5 |  |  |  |
| Cingulate Gyrus, posterior division |  | 13.3 |  |  |  |
| Precuneous Cortex |  | 18.3 |  |  |  |
| Cluster #14 |  | Average Atlas Probability Within Cluster |  |  |  |
| Temporal Pole |  | 7.0 |  |  |  |
| Superior Temporal Gyrus, posterior division |  | 8.2 |  |  |  |
| Middle Temporal Gyrus, anterior division |  | 7.2 |  |  |  |
| Middle Temporal Gyrus, posterior division |  | 23.9 |  |  |  |
| Middle Temporal Gyrus, temporooccipital part |  | 4.7 |  |  |  |
| Cluster #13 |  | Average Atlas Probability Within Cluster |  |  |  |
| Left Crus I |  | 25.6 |  |  |  |
| Left Crus II |  | 46.9 |  |  |  |
| Cluster #12 |  | Average Atlas Probability Within Cluster |  |  |  |
| Pre-Supplementary Motor Area |  | 6.6 |  |  |  |
| Area 9 |  | 7.9 |  |  |  |
| Area 8B |  | 10.0 |  |  |  |
| Cluster #11 |  | Average Atlas Probability Within Cluster |  |  |  |
| Angular Gyrus |  | 28.5 |  |  |  |
| Lateral Occipital Cortex, superior division |  | 29.1 |  |  |  |
| Cluster #10 |  | Average Atlas Probability Within Cluster |  |  |  |
| Right Crus I |  | 48.9 |  |  |  |
| Right Crus II |  | 33.4 |  |  |  |
| Cluster #9 |  | Average Atlas Probability Within Cluster |  |  |  |
| IFS (Inferior Frontal Sulcus) |  | 7.6 |  |  |  |
| FPI (Lateral Frontal Pole) |  | 11.1 |  |  |  |
| FPM (Medial Frontal Pole) |  | 6.1 |  |  |  |
| Cluster #8 |  | Average Atlas Probability Within Cluster |  |  |  |
| Area 47 |  | 16.9 |  |  |  |
| FPI (Lateral Frontal Pole) |  | 6.7 |  |  |  |
| Cluster #7 |  | Average Atlas Probability Within Cluster |  |  |  |
| Superior Temporal Gyrus, posterior division |  | 10.1 |  |  |  |
| Middle Temporal Gyrus, posterior division |  | 47.5 |  |  |  |
| Middle Temporal Gyrus, temporooccipital part |  | 5.3 |  |  |  |

**Table 4.** Atlas query of conjunction of Right Crus II connectivity results. For large clusters, sub-region labels are listed below. COG: Center of Gravity.

| ClusterIndex | Voxels | COG X (mm) | COG Y (mm) | COG Z (mm) | COG Label |
| --- | --- | --- | --- | --- | --- |
| 16 | 2775 | 28.1 | -73.8 | -40.5 | 96% Right Crus II |
| 15 | 2573 | -3.58 | -49.1 | 31.6 | 76% Cingulate Gyrus |
| 14 | 1901 | -40.5 | -69.7 | 36.9 | 51% Lateral Occipital Cortex, superior division |
| 13 | 1873 | -2.94 | 59.3 | -0.838 | 41% Area 10 |
| 12 | 1183 | -60.2 | -48.6 | -10.1 | 50% Middle Temporal Gyrus, temporooccipital part |
| 11 | 982 | -22.8 | 27.7 | 48.3 | 58% Area 8B |
| 10 | 777 | -59.3 | -12 | -15.4 | 33% Middle Temporal Gyrus, posterior division |
| 9 | 484 | 3.45 | -53.6 | -48.1 | 95% Right IX |
| 8 | 209 | -37.7 | 35.1 | -12.6 | 34% Frontal Orbital Cortex |
| 7 | 163 | -13.1 | -87.5 | -38.7 | 99% Left Crus II |
| 6 | 44 | -26.7 | -17.1 | -14.5 | 78% Left Hippocampus |
| 5 | 27 | -45.2 | 50.1 | 0.148 | 100% Area 46 |
| ----- |  |  |  |  |  |
| Structures to which each cluster belongs to: |  |  |  |  |  |
| Cluster #16 |  |  | Average Atlas Probability Within Cluster |  |  |
| Right Crus I |  |  | 18.4 |  |  |
| Right Crus II |  |  | 50.4 |  |  |
| Cluster #15 |  |  | Average Atlas Probability Within Cluster |  |  |
| Cingulate Gyrus, posterior division |  |  | 31.6 |  |  |
| Precuneous Cortex |  |  | 33.5 |  |  |
| Cluster #13 |  |  | Average Atlas Probability Within Cluster |  |  |
| Area 10 |  |  | 23.5 |  |  |
| Frontal Medial Cortex |  |  | 14.7 |  |  |
| Paracingulate Gyrus |  |  | 7.0 |  |  |
| Cluster #12 |  |  | Average Atlas Probability Within Cluster |  |  |
| Middle Temporal Gyrus, posterior division |  |  | 12.3 |  |  |
| Middle Temporal Gyrus, temporooccipital part |  |  | 22.9 |  |  |
| Inferior Temporal Gyrus, temporooccipital part |  |  | 14.4 |  |  |
| Cluster #11 |  |  | Average Atlas Probability Within Cluster |  |  |
| Cluster2 (PreSMA) |  |  | 12.4 |  |  |
| Cluster10 (area 8B) |  |  | 31.9 |  |  |
| Cluster #10 |  |  | Average Atlas Probability Within Cluster |  |  |
| Superior Temporal Gyrus, posterior division |  |  | 9.4 |  |  |
| Middle Temporal Gyrus, anterior division |  |  | 19.2 |  |  |
| Middle Temporal Gyrus, posterior division |  |  | 25.4 |  |  |
| Cluster #8 |  |  | Average Atlas Probability Within Cluster |  |  |
| Frontal Pole |  |  | 27.0 |  |  |
| Frontal Orbital Cortex |  |  | 33.3 |  |  |

**Table 5.**
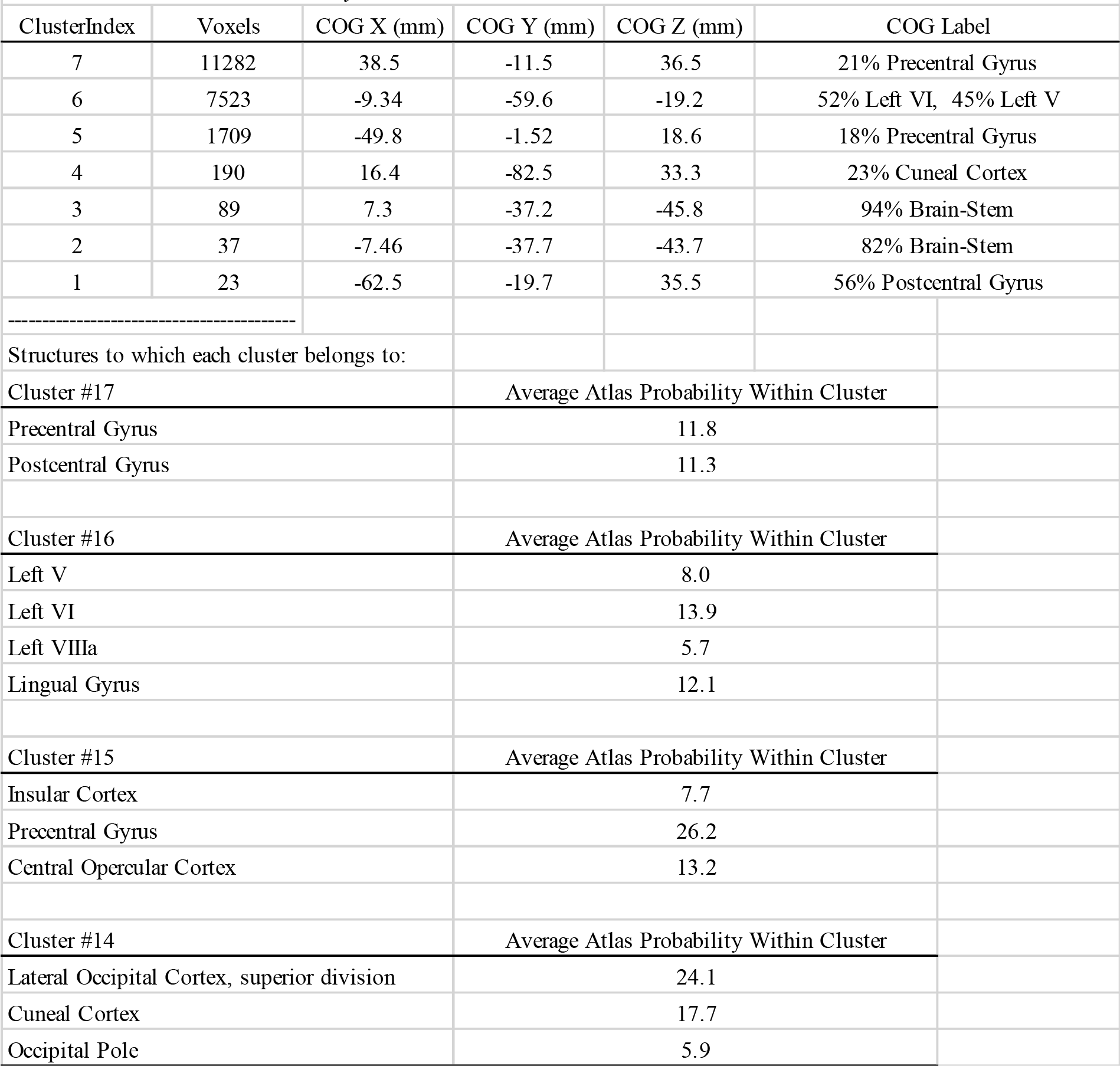
Atlas query of conjunction of Left Lobule VI connectivity results. For large clusters, sub-region labels are listed below. COG: Center of Gravity.

**Table 6.**
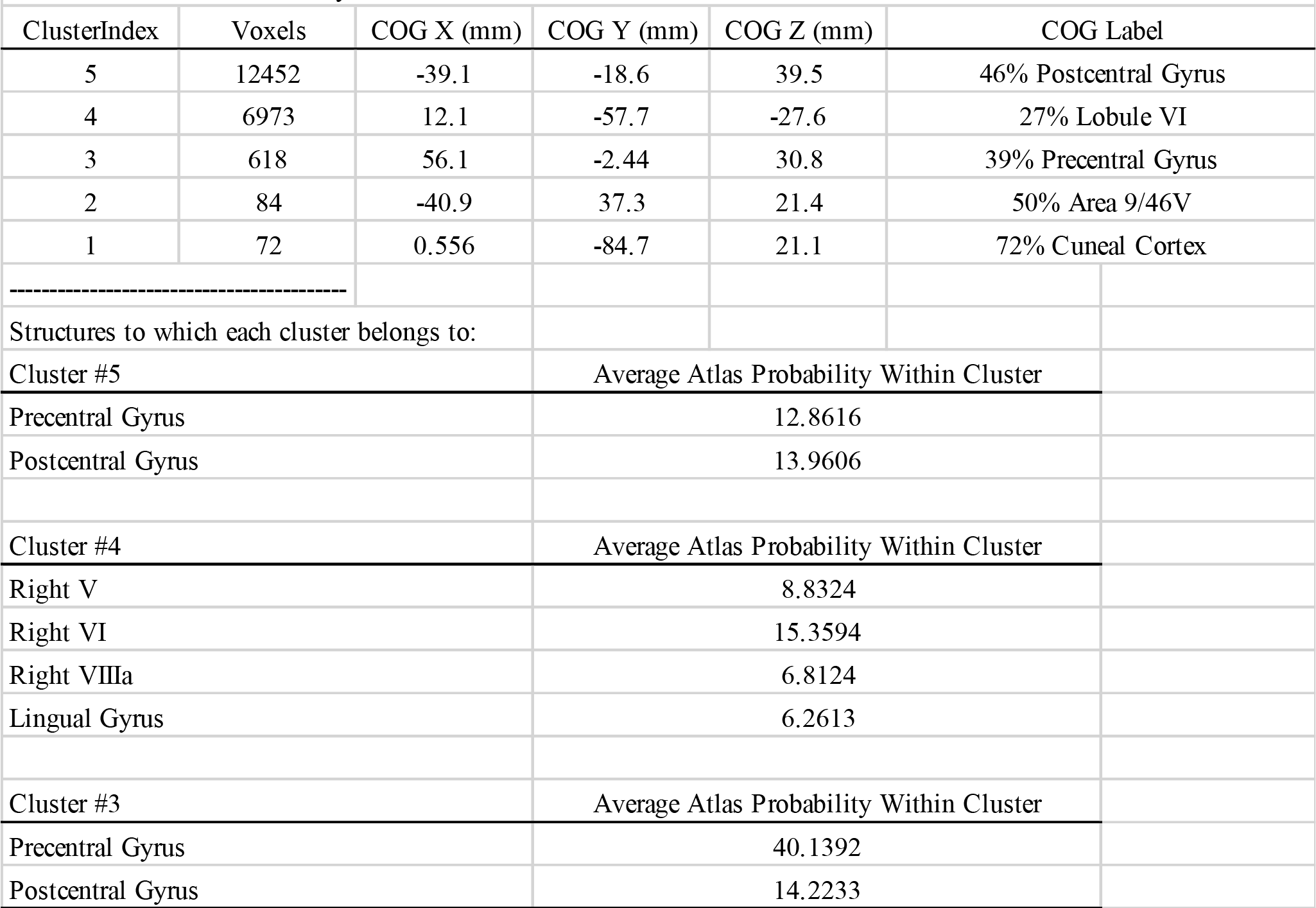
Atlas query of conjunction of Right Crus II connectivity results. For large clusters, sub-region labels are listed below. COG: Center of Gravity.

Examining Figure 1, the prefrontal connectivity was not as clear for the Right Crus II conjunction compared to the Left Crus II conjunction. Therefore, we examined the direct comparisons of Crus II to Crus I and Lobule VI to help clarify the patterns of connectivity. As shown in Figure 2, Left Crus II showed stronger connectivity compared to Crus I and Lobule VI in bilateral posterior DLPFC, bilateral anterior ventrolateral PFC, and medial area 9 (see also Supplemental Tables 5 & 7). When examining the connectivity of Right Crus II, Figure 3 illustrates that the comparison of Right Crus II and Lobule VI shows similar patterns of connectivity as the Left Crus II conjunction, but Right Crus II showed greater connectivity compared to Right Crus I in a cluster in the middle DLPFC, in between the posterior DLPFC and anterior ventrolateral PFC clusters (see also Supplemental Tables 6 & 8). This middle DLPFC cluster more closely resembles the predicted region of connectivity with Crus II, extending the length of ventral 9/46 [Petrides and Pandya, 1994], a region implicated in working memory function in humans and macaques [Petrides, 2005].

#### Lobule VI

As shown in Figure 1, there was little prefrontal connectivity with Left Lobule VI, with the bulk of connectivity located in the sensorimotor cortex. The conjunction for Right Lobule VI, however, showed a small cluster of connectivity in the left DLPFC. Turning to the direct comparison of Lobule VI and Crus I, there was no prefrontal connectivity for Left Lobule VI, but Right Lobule VI was more connected with the left DLPFC than Right Crus I (See Figures 2&3, Supplemental Tables 9-12). Comparing Lobule VI and Crus II, for the left and right ROIs there was connectivity with bilateral anterior DLPFC (areas 46 and dorsal 9/46). This area of connectivity was slightly caudal to the RLPFC cluster connected with Crus I. This finding was somewhat surprising, and contrary to our predictions, as Lobule VI connectivity patterns at rest have implicated premotor cortical regions [Bernard et al., 2012]. However, task-based activation patterns do suggest contributions to higher-order processing [E et al., 2012; Guell et al., 2018b; Stoodley et al., 2012; Stoodley and Schmahmann, 2009]. Further, in their investigation of cerebellar sensorimotor networks, Kipping and colleagues [2013] demonstrated a similar pattern of connectivity, implicating the lateral prefrontal cortex.

#### Functional Connectivity Interim Summary

Comparisons across Crus I, Crus II, and Lobule VI suggest that their patterns of connectivity vary across the prefrontal cortex in a manner suggesting parallel PFC and cerebellar functional processing gradients. Crus I is most strongly associated with prefrontal cortical regions associated with higher level control, while Crus II was associated with areas implicated in working memory and more specific cognitive processes (as opposed to more abstract processing). Finally, Lobule VI was also correlated with many of these working memory regions, and somewhat surprisingly, there was no unique connectivity with areas associated with the planning of motoric responses, counter to our predictions.

### Task-Based Functional Activation Results

#### Relational Processing Task

For the relational processing task, we examined activation for the Relational vs. Match contrast. This task has been suggested to serve as a localizer for RLPFC/ area 10 [Smith et al., 2007]. While we did observe rostral prefrontal activation, the activation was ventral to the expected area 10, in anterior 47r (See Figure 4 and Table 7). This cluster is similar to that which showed connectivity with Crus II, further differentiating these findings from our predictions. Additional prefrontal activation was found in the anterior insula, posterior DLPFC, dorsal medial prefrontal cortex (area 8), and dorsal premotor cortex. In the cerebellum, there was activation of both Crus I and Crus II, as well as Lobule VI and Lobule VIIb. Smith and colleagues [2007] demonstrated that the Relational Processing Task robustly activates the RLPFC (i.e., the lateral portion of BA10) across participants, however, they did not report cluster coordinates for the group average, only reporting activation foci in BA10 for each participant. Nevertheless, for most participants, activation foci were located in the Superior and Middle Frontal Gyri, while area 47 is located in the Inferior Frontal Gyrus. In their multimodal cortical parcellation using the HCP data, Glasser and colleagues also found that area 47r was active in the Relational vs. Match contrast [Glasser et al., 2016; see Figure 23 of their Supplementary Neuroanatomical Results].

**Figure 4.**
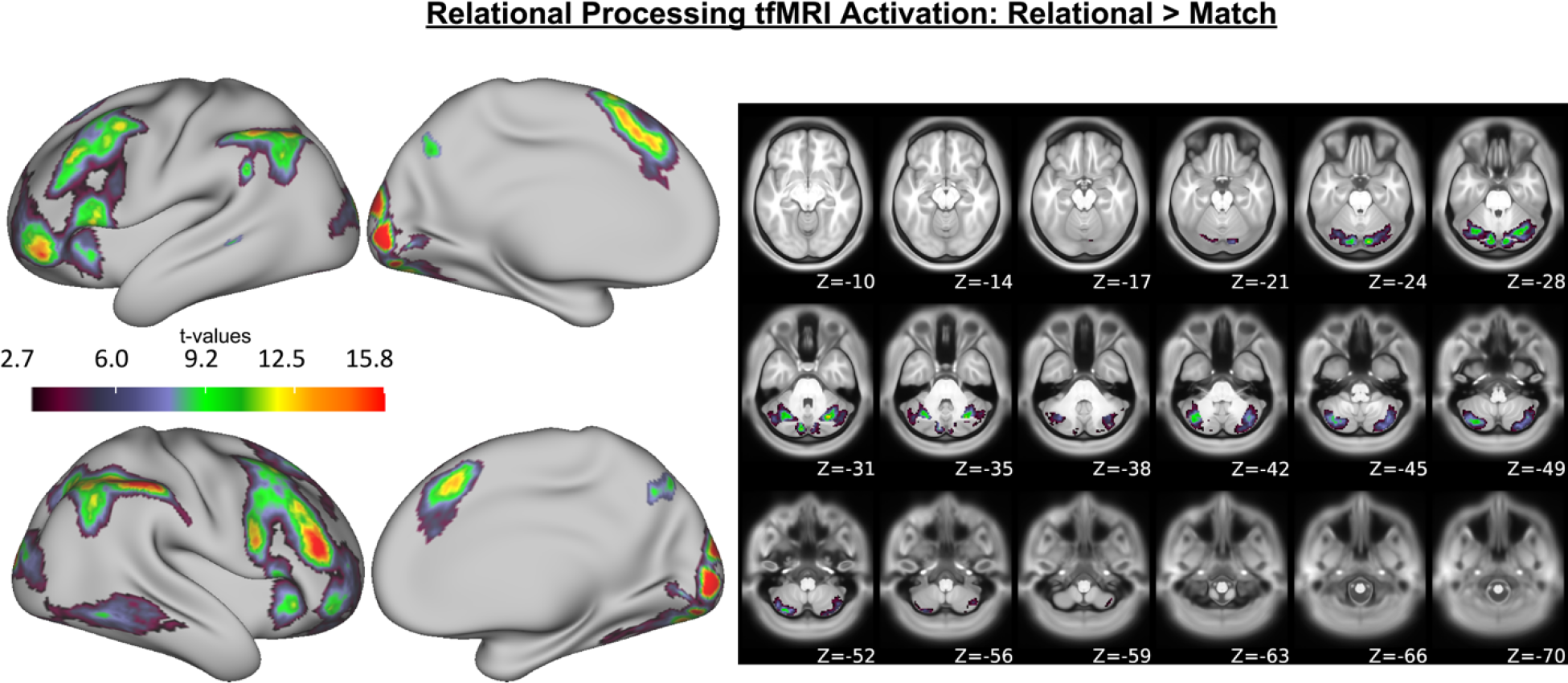
Relational processing related cortical surface activation (left) and cerebellar volumetric activation (right). Results come from a group-level analysis of the contrast of relational vs. match for the Relational Processing Task.

**Table 7.** Areas of cortical surface activation during the relational processing task.

| *X* | *Y* | *Z* | *Glasser Region* | *X* | *Y* | *Z* | *Glasser Region* |
| --- | --- | --- | --- | --- | --- | --- | --- |
| -43.16 | 51.75 | -10.99 | Left Area anterior 47r | 46.72 | 44.85 | 20.86 | Right Area posterior 9/46v |
| -45.71 | 51.13 | -2.86 | Left Area posterior 47r | 38.97 | 30.76 | -8.41 | Right Anterior Ventral Insular Area |
| -51.39 | 26.05 | 5.63 | Left Area 44 | 43.24 | 51.89 | -12.2 | Right anterior Area 47r |
| -42.94 | 34.55 | 39.16 | Left Area 8C | 45.57 | 52.54 | -6.7 | Right posterior Area 47r |
| -44.74 | 18.57 | 45.49 | Left Area 8C | 56.65 | 14.82 | 21.17 | Right Rostral Area 6 |
| -44.52 | 11.15 | 52.84 | Left Area 55b | 44.02 | 35.68 | 39.17 | Right Area 8C |
| -36.33 | 24.07 | 55.72 | Left Ventral Area 8A | 46.16 | 18.2 | 42.59 | Right Area 8C |
| -11.94 | 16.88 | 66.78 | Left Superior Frontal Language Area | 38.57 | 24.27 | 55.31 | Right Area 8Av |
| -5.96 | 28.69 | 48.5 | Left Area 8B medial | 36.42 | 16.7 | 59.66 | Right Inferior 6-9 Transitional Area |
| -51.51 | -52.34 | 49.8 | Left Area PFm Complex | 6.6 | 29.18 | 48.98 | Right Area 8B medial |
| -42.38 | -68.67 | 49.15 | Left Area PFm Complex | 59.89 | -33.48 | 46.96 | Right Area PF Complex |
| -7.77 | -72.25 | 41.87 | Left Parieto-Occipital Sulcus Area 2 | 52.36 | -39.92 | 50.31 | Right Area IntraParietal 2 |
| -15.61 | -90.05 | -13.56 | Left Second Visual Area | 55.99 | -41.9 | 48.7 | Right Area PFm Complex |
| -28.53 | -83.15 | -17.26 | Left Fourth Visual Area | 45.79 | -52.63 | 50.2 | Right Area IntraParietal 2 |
| -10.04 | -95.63 | 0 | Left Primary Visual Cortex | 46.83 | -63.46 | 48.24 | Right Area PFm Complex |
| -10.63 | -97.52 | 13.78 | Left Second Visual Area | 40.08 | -68.04 | 48.98 | Right Area IntraParietal 1 |
| -19.14 | -100.1 | 14.97 | Left Third Visual Area | 29.31 | -72.5 | 49.12 | Right Medial IntraParietal Area |
| - | - | - | - | 32 | -78.71 | -17.04 | Right Eighth Visual Area |
| - | - | - | - | 24.99 | -82.21 | -12.27 | Right Fourth Visual Area |
| - | - | - | - | 19.1 | -88.21 | -12.27 | Right Second Visual Area |
| - | - | - | - | 21.72 | -86.92 | -13.33 | Right Third Visual Area |
| - | - | - | - | 13.74 | -94.47 | -0.07 | Right Primary Visual Cortex |
| - | - | - | - | 14.69 | -96.49 | 15.02 | Right Second Visual Area |
| - | - | - | - | 20.43 | -97.23 | 18.69 | Right Third Visual Area |

*Note:* XYZ Coordinates are reported in inflated cortical surface space. More information about Glasser Parcellation areas can be found in the supplementary neuroanatomical results of Glasser et al. (2016).

#### Working Memory Task

To assess brain activity associated with working memory, we used the 2-back vs. 0-back contrast from the embedded n-back task. Both the RLPFC and DLPFC were activated in this contrast, as shown in Figure 5 (see also Table 8). The RLPFC activity was located in left area 9/46d and right area 10p. This area of activation was similar to the location that showed functional connectivity with Crus I. As shown in Figure 8, the DLPFC activation overlapped with the activation from the Relational Processing Task, suggesting this region serves a similar purpose in both tasks, likely working memory. There was also prefrontal activation in the dorsal premotor cortex and dorsal medial prefrontal cortex, largely overlapping with the Relational Processing Task activation. In the cerebellum, there was widespread activation, with Crus I and Crus II both activated, as well as Lobules I-V, V, VIIIb, and IX.

**Figure 5.**
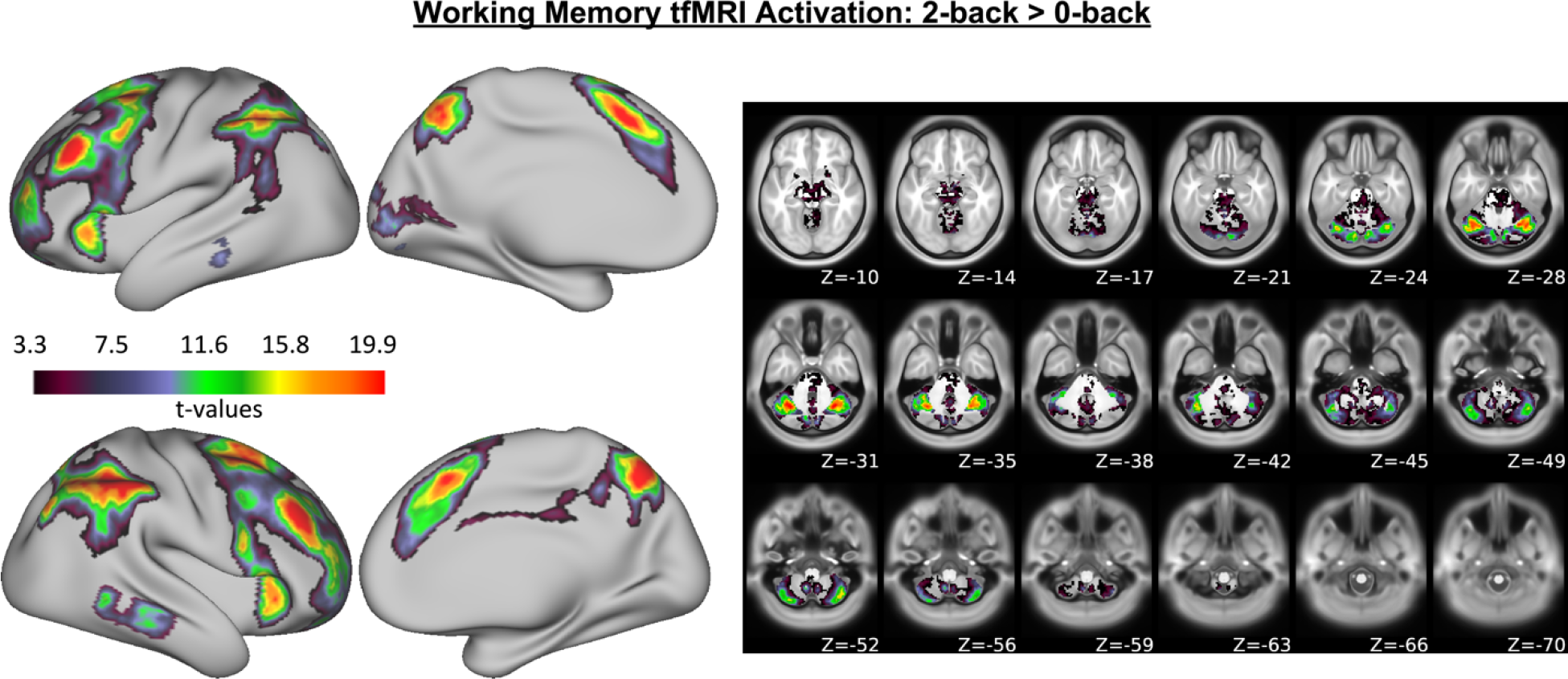
Working memory related cortical surface activation (left) and cerebellar volumetric activation (right). Results come from a group-level analysis of the contrast of 2-back vs. 0-back trials for the N-Back Task.

**Table 8.**
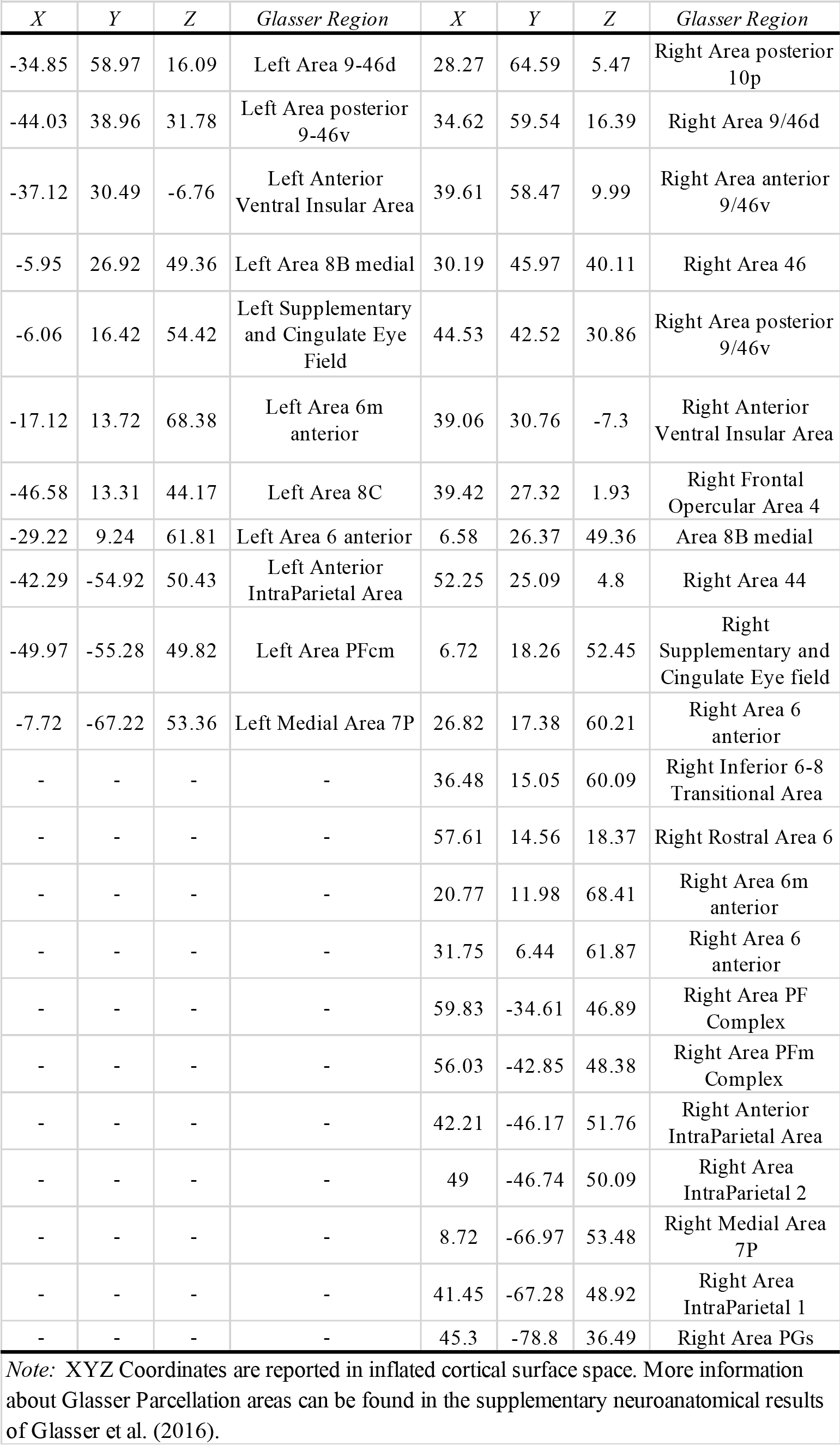
Areas of cortical surface activation during the working memory task

| *X* | *Y* | *Z* | *Glasser Region* | *X* | *Y* | *Z* | *Glasser Region* |
| --- | --- | --- | --- | --- | --- | --- | --- |
| -34.85 | 58.97 | 16.09 | Left Area 9-46d | 28.27 | 64.59 | 5.47 | Right Area posterior 10p |
| -44.03 | 38.96 | 31.78 | Left Area posterior 9-46v | 34.62 | 59.54 | 16.39 | Right Area 9/46d |
| -37.12 | 30.49 | -6.76 | Left Anterior Ventral Insular Area | 39.61 | 58.47 | 9.99 | Right Area anterior 9/46v |
| -5.95 | 26.92 | 49.36 | Left Area 8B medial | 30.19 | 45.97 | 40.11 | Right Area 46 |
| -6.06 | 16.42 | 54.42 | Left Supplementary and Cingulate Eye Field | 44.53 | 42.52 | 30.86 | Right Area posterior 9/46v |
| -17.12 | 13.72 | 68.38 | Left Area 6m anterior | 39.06 | 30.76 | -7.3 | Right Anterior Ventral Insular Area |
| -46.58 | 13.31 | 44.17 | Left Area 8C | 39.42 | 27.32 | 1.93 | Right Frontal Opercular Area 4 |
| -29.22 | 9.24 | 61.81 | Left Area 6 anterior | 6.58 | 26.37 | 49.36 | Area 8B medial |
| -42.29 | -54.92 | 50.43 | Left Anterior IntraParietal Area | 52.25 | 25.09 | 4.8 | Right Area 44 |
| -49.97 | -55.28 | 49.82 | Left Area PFcm | 6.72 | 18.26 | 52.45 | Right Supplementary and Cingulate Eye field |
| -7.72 | -67.22 | 53.36 | Left Medial Area 7P | 26.82 | 17.38 | 60.21 | Right Area 6 anterior |
| - | - | - | - | 36.48 | 15.05 | 60.09 | Right Inferior 6-8 Transitional Area |
| - | - | - | - | 57.61 | 14.56 | 18.37 | Right Rostral Area 6 |
| - | - | - | - | 20.77 | 11.98 | 68.41 | Right Area 6m anterior |
| - | - | - | - | 31.75 | 6.44 | 61.87 | Right Area 6 anterior |
| - | - | - | - | 59.83 | -34.61 | 46.89 | Right Area PF Complex |
| - | - | - | - | 56.03 | -42.85 | 48.38 | Right Area PFm Complex |
| - | - | - | - | 42.21 | -46.17 | 51.76 | Right Anterior IntraParietal Area |
| - | - | - | - | 49 | -46.74 | 50.09 | Right Area IntraParietal 2 |
| - | - | - | - | 8.72 | -66.97 | 53.48 | Right Medial Area 7P |
| - | - | - | - | 41.45 | -67.28 | 48.92 | Right Area IntraParietal 1 |
| - | - | - | - | 45.3 | -78.8 | 36.49 | Right Area PGs |

*Note:* XYZ Coordinates are reported in inflated cortical surface space. More information about Glasser Parcellation areas can be found in the supplementary neuroanatomical results of Glasser et al. (2016).

#### Language Task

To assess language processing, we used the Story vs. Math contrast. This contrast activated both the left and right posterior inferior frontal cortex (areas 45 and 47 lateral), the medial frontal pole, ventral area 8A, and the left Superior Frontal Language Area (see Figure 6 and Table 9). Unlike the other 3 tasks, language activated large areas of the temporal cortex, specifically, bilateral anterior temporal cortex and temporal parietal junction. In the cerebellum, bilateral Crus II and Lobule XI were activated, with the Crus II peak activation being more lateral to that of the Relational Processing Task, as shown in Figure 8.

**Figure 6.**
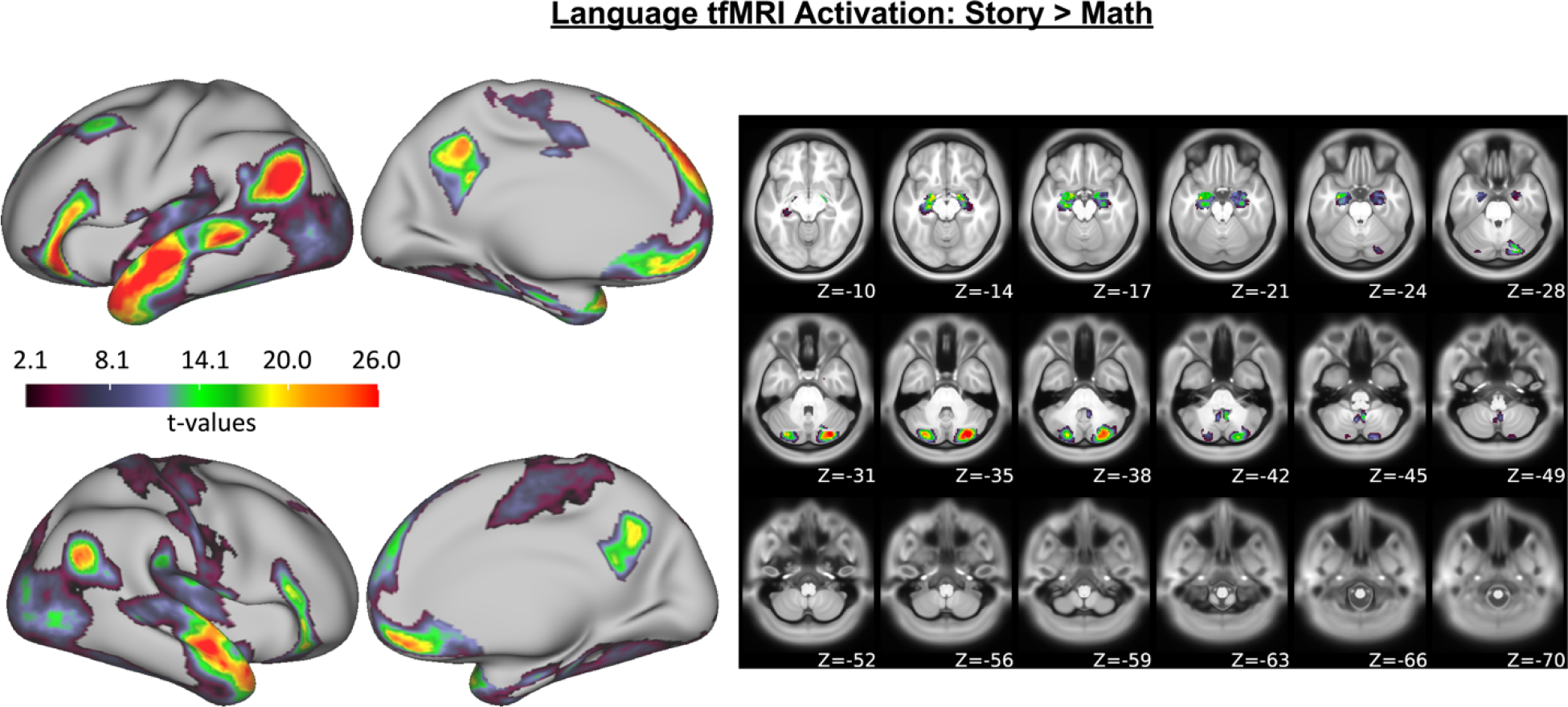
Language related cortical surface activation (left) and cerebellar volumetric activation (right). Results come from a group-level analysis of the contrast of story vs. math trials in the Story Task.

**Table 9.**
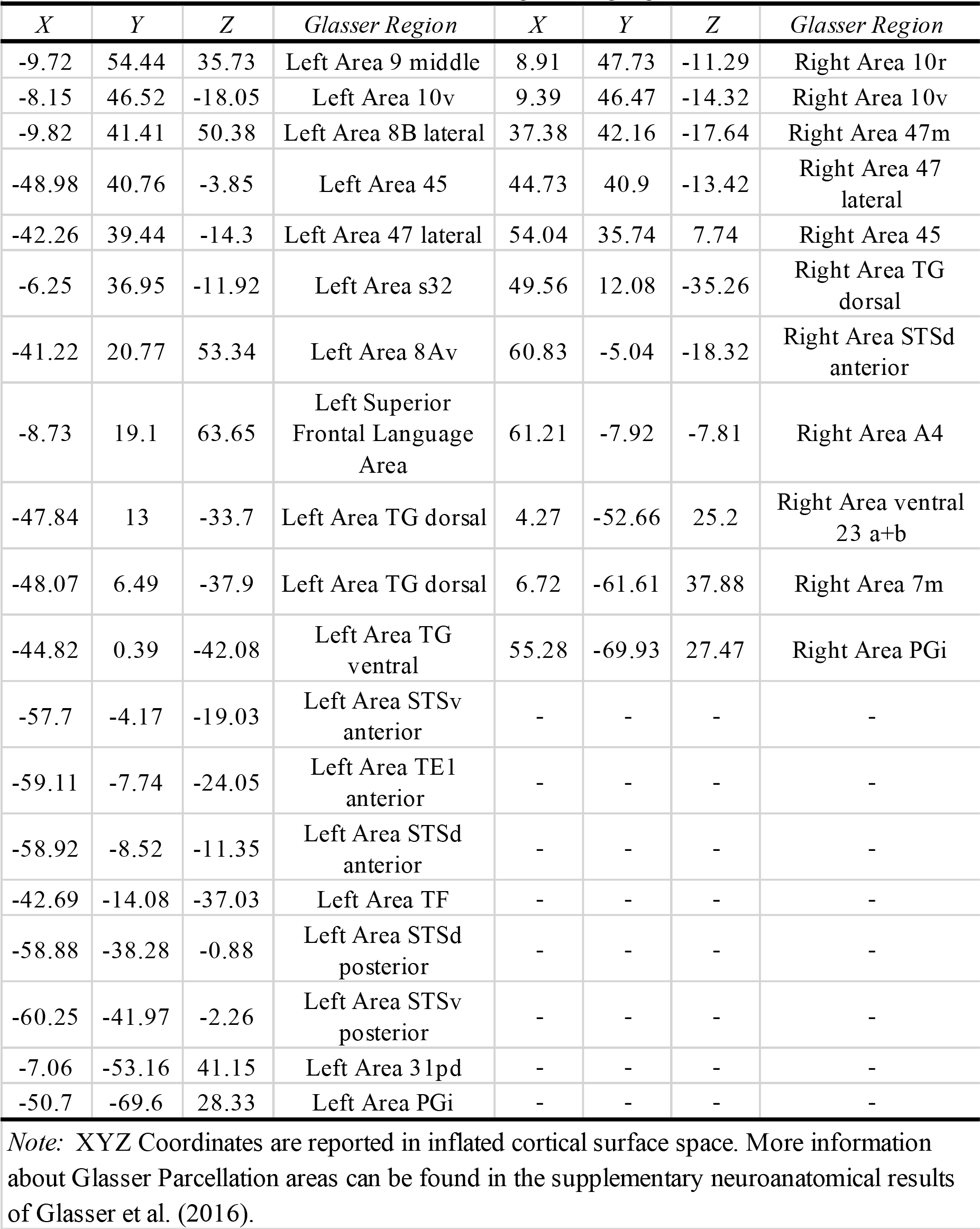
Areas of cortical surface activation during the language task.

#### Motor Task

To assess motor function, we used the Right Hand contrast. Unsurprisingly, this was associated with left sensorimotor cortex and premotor cortex, as shown in Figure 7 and Table 10. In the cerebellum, Right Lobules VIIIb and V were activated.

**Figure 7.**
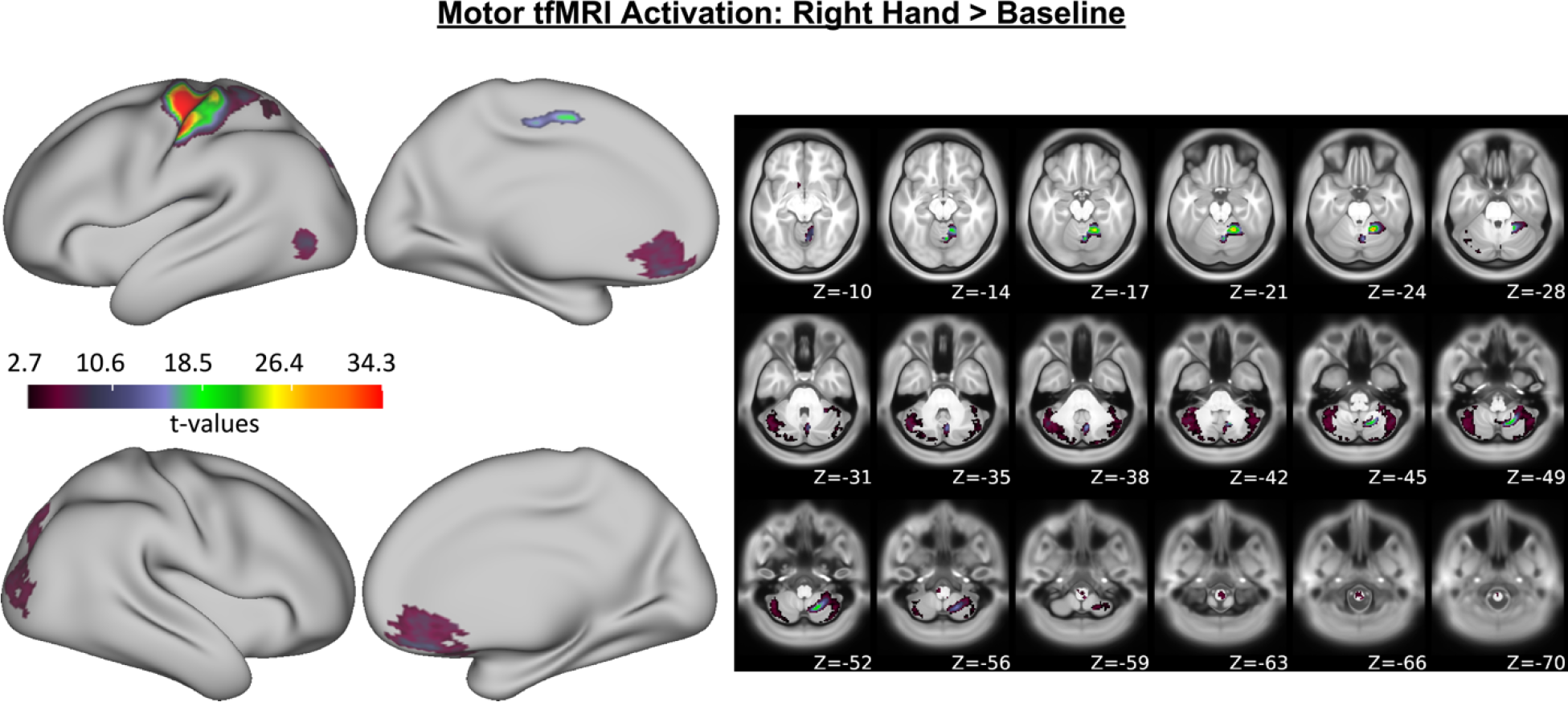
Motor related cortical surface activation (left) and cerebellar volumetric activation (right). Results come from a group-level analysis of the contrast of right hand vs. baseline trials in the Motor Tapping Task.

**Table 10.** Areas of cortical surface activation during the motor task.

| *X* | *Y* | *Z* | *Glasser Region* |
| --- | --- | --- | --- |
| -6.01 | -7.31 | 55.95 | Left Dorsal Area 24d |
| -35.6 | -14.14 | 68.77 | Left Dorsal Area 6 |
| -37.86 | -18.7 | 65.41 | Left Area 4 |
| -48.39 | -19.02 | 54.7 | Left Primary Sensory Cortex |
| -54.06 | -19.48 | 52.11 | Left Area 1 |
| -37.86 | -24.06 | 62.07 | Left Area 3a |
| -40.81 | -33.98 | 66.9 | Left Area 1 |

*Note:* XYZ Coordinates are reported in inflated cortical surface space. More information about Glasser Parcellation areas can be found in the supplementary neuroanatomical results of Glasser et al. (2016).

#### Task-Related Activation Interim Summary

The analysis of task-related functional activation provided further evidence for existence of functional gradients of CB-PFC networks. As shown in Figure 8, there were dissociable areas of activation for all four tasks in the prefrontal cortex and cerebellum. There were overlapping areas of activation for the relational processing task and working memory task in the mid- and posterior DLPFC. In the cerebellum, these tasks showed overlapping areas of activation in both Crus I and Crus II, but the activation for the relational processing task was more posterior to that of the working memory task. However, there were some notable deviations from our predictions. As shown in Figure 8, in the RLPFC, the activation for the relational processing task and working memory task were reversed from what we predicted, with relational processing being more ventral, in a region connected more with Crus II than Crus I (see Figure 1). In addition, the relational processing task activated Crus II to a greater extent than the working memory task.

**Figure 8.**
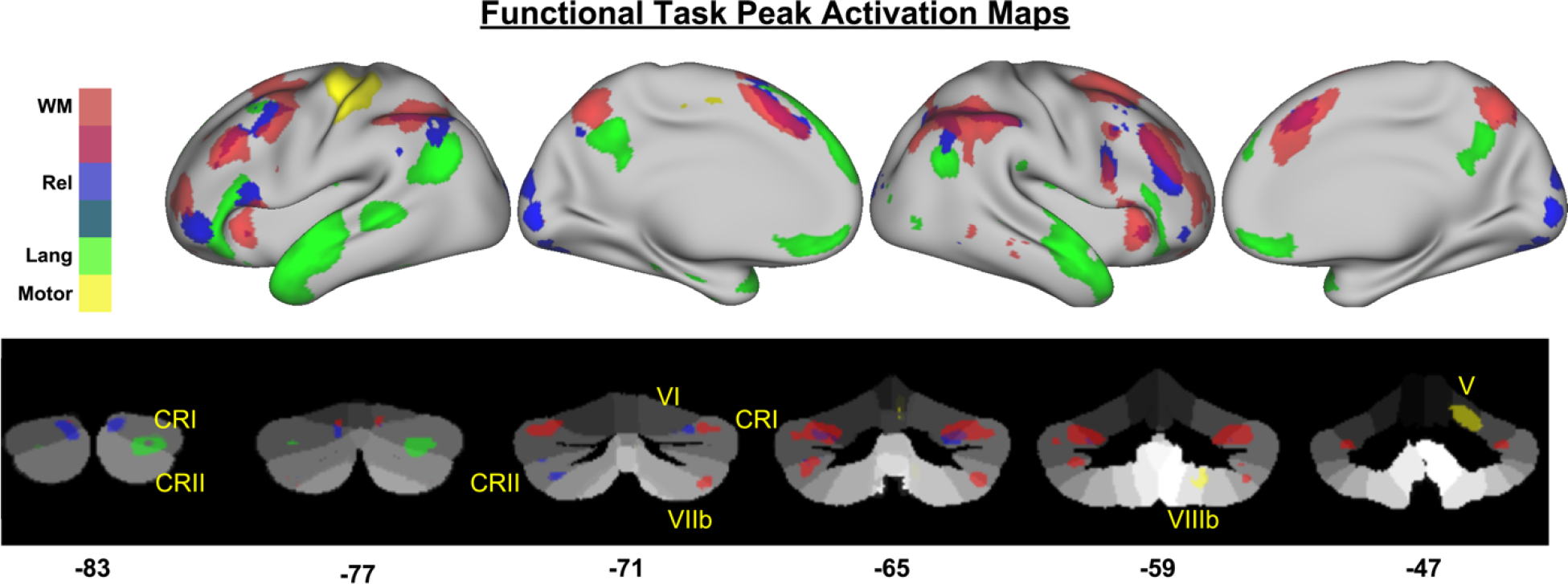
Peak activations from the four functional tasks: working memory (red), relational processing (blue), language (green), and motor (yellow) overlaid on the inflated cortical surface (top row in each section) and the cerebellum (bottom row in each section). Cluster-corrected activation maps were further thresholded at the 95 percentile to isolate peaks of activation, and were binarized for display. The working memory activation was thresholded at T > 10.98, relational processing was thresholded at T > 9.34, language was thresholded at T > 15.5, and motor was thresholded at T > 13.45.

## Discussion

The goal of this study was to test the hypothesis that cerebellar-prefrontal connections are organized along a gradient. Across several domains, it has been demonstrated that the lateral prefrontal cortex is organized along a rostral-caudal gradient of abstraction, with anterior PFC associated with the most abstract levels of processing (e.g., long-term goals and integrating information across time levels) and posterior PFC associated with concrete information such as stimuli and responses. While previous work has demonstrated the possibility of a similar gradient in the basal ganglia [Nee and Brown, 2013; Verstynen et al., 2012], it is unclear if the cerebellum follows a similar gradient. Previous work has suggested that there is a functional topography for nonmotor tasks in the cerebellum [E et al., 2012; Guell et al., 2018a; Guell et al., 2018b; Stoodley et al., 2012], yet, it is unclear whether the functional topography in the cerebellum shows an abstraction gradient.

While the current study was limited in that we relied on tfMRI from tasks not necessarily optimized to investigate this purported functional gradient, the findings suggest that such a gradient of abstraction may be present in the cerebellum. There is limited evidence from task-based fMRI to support this hypothesis, however. To our knowledge, previous studies of levels of abstraction in EF have not focused on the cerebellum. The cerebellum is frequently not imaged due to limitations in coverage with standard imaging protocols. Moreover, standard spatial smoothing kernels employed in most cortical/whole-brain fMRI preprocessing pipelines (e.g., 6-8 mm FWHM) may be too large to allow for the separation of neighboring lobules [Bernard et al., 2012; Bernard and Seidler, 2013]. Nevertheless, the suggestion that such a functional gradient exists in the cerebellum is supported by a number of previous studies [e.g., Balsters et al., 2013; Stoodley et al., 2012; Stoodley and Schmahmann, 2009].

We found that Crus I was connected to the RLPFC more strongly than Crus II and Lobule VI. This is in line with our prior work which investigated the connectivity of each cerebellar lobule, but did not compare the networks of different lobules [Bernard et al., 2012]. However, there was mixed evidence for our prediction that the Relational Processing Task would be associated with activation in both Crus I and RLPFC. While there was anterior PFC activation for the task, it was more ventral than the region typically referred to as RLPFC. Nevertheless, the Relational Processing Task did show activation in Crus I. There is evidence to suggest that Crus I is involved in higher-order cognitive processing, in concert with anterior prefrontal cortex. We have previously shown that Crus I and the rostral lateral prefrontal cortex are activated when participants perform voluntary task switching, a task that requires them to select tasks according to abstract, internally maintained task goals [Orr and Banich, 2014]. The rostral lateral prefrontal cortex has also been linked to prospective memory, i.e., remembering to perform an action after a delay [Burgess et al., 2003]. Several studies of prospective memory have shown activation of Crus I [Burgess et al., 2003; Reynolds et al., 2009]. Further, when examining individuals who don’t show multitasking costs, aka “supertaskers”, Medeiros-Ward and colleagues [2014] found that Crus I showed a group-by-load effect along with the rostral lateral prefrontal cortex.

While a number of studies have implicated a role of Crus I in executive function, there is less evidence for a role of Crus II. In an fMRI meta-analysis, Stoodley & Schmahmann [2009] found strong activation overlap in Crus II during language tasks. Similarly, here, in our own tfMRI analysis of language processing using HCP data, we also found cerebellar activation in Crus II. The current connectivity findings also support a link between Crus II and language, as Crus II was connected with language-related regions of posterior inferior frontal gyrus, including areas 45 and 47. However, Buckner and colleagues [2011] identified a connection between Crus II and the default mode network. If the center of gravity of the Right Crus II seed (from the SUIT atlas thresholded at 50%, coordinates: 26, −76, −42) is entered into *neurosynth* [Yarkoni et al., 2011], top meta-analytic terms associated with the region include “past”, “socially”, and “autobiographical”. Nevertheless, the functional connectivity and meta-analytic connectivity maps for the same region from *neurosynth* (http://neurosynth.org/locations/?y=-76&x=26&z=-42) contains left posterior inferior frontal gyrus cortex. Thus, while several lines of evidence support a role for Crus II in language, further studies are needed to clarify the nature of this role.

While we predicted that Lobule VI would should show connectivity with posterior prefrontal cortex, the connectivity was more posterior in primary motor cortex, rather than premotor cortex. Prior studies have demonstrated connectivity between Lobule VI and premotor cortex [Bernard et al., 2012], though Kipping and colleagues [2013] demonstrated associations with more lateral prefronal cortical regions. The conflicting results here may however be due to the nature of our analyses where we directly compared Lobule VI with Crus I and II. As seen in Figure 2, both Crus I and II also have areas of connectivity with dorsal and ventral pre-motor cortical regions, and thus it may be the case that the greater connectivity for Lobule VI is with more primary motor cortical regions. However, it is also notable that the sample here is at much higher resolution and has a significantly larger sample than prior work [Bernard et al., 2012]. Reineberg and Banich [2016] showed that individual difference in network dynamics at rest in Lobule VI were associated with working memory updating. Further, meta-analytic evidence suggests that this region is involved not only in working memory, but across motor tasks and learning, and in language as well [E et al., 2012; Stoodley & Schmahmann, 2009; Bernard & Seidler, 2013]. The seemingly diverse roles and contributions of Lobule VI suggest that it is a transition area of sorts such that there is involvement in more abstract higher order thinking to some degree, but also makes contributions to motor planning. Because much of the evidence in this regard comes from meta-analysis, it is not feasible to dissociate the more concrete motor-reliant execution aspects of these tasks from the more abstract processing. However, the stronger resting state associations with motor cortical regions that that we see with Lobule VI, particularly relative to Crus I and Crus II, suggest that perhaps the role of this cerebellar region is with respect to the more concrete aspects of motor response selection during the performance of higher order cognitive tasks. Future work specifically dissociating these levels of abstraction with respect to the cerebellum is clearly warranted in the future.

Here, using a large sample of individuals from the HCP, we carefully probed both the functional connectivity and activation of cerebellar lobules with respect to one another, as a first step towards understanding cerebellar contributions to executive function. While past work has taken a targeted lobular approach [Bernard et al., 2012], looked at the cerebellum more generally [Habas and Cabanis, 2007; O’Reilly et al., 2010], or made comparisons framed by the prefrontal cortex broadly defined [Krienen & Buckner, 2009], this cerebellar approach allows us to carefully interrogate connectivity patterns that are of potential importance to our understanding of EF. Broadly speaking, our results suggest that a gradient of abstraction may also be present in the cerebellum, paralleling what is see in the PFC [e.g., Badre, 2008], likely subserved by the closed-loop circuitry linking these two disparate brain regions [Bernard et al., 2016; Kelly and Strick, 2003; Salmi et al., 2010; Strick et al., 2009]. While the exact role of the cerebellum in non-motor behavior remains unknown, it has been suggested that the structure acts to process internal models of thought [Ramnani, 2006; Ito, 2008; Ramnani, 2014], much as is done in the motor domain [e.g., Imamizu et al., 2000]. Across cerebellar lobules, the cytoarchitectonics remain the same, only the cortical connections change, suggesting a similar computation is being performed, just on distinct inputs [Ramnani, 2006; Ito, 2008; Ramnani, 2014]. In his more recent writings on this topic, Ramnani [2014] has suggested that the cerebellum supports more automated processes after learning, as compared to the more cognitively demanding top-down processes that occur early on when performing a task. With damage and disease, one would therefore experience deficits in thought and processing, such as those seen in patients with schizophrenia [Andreasen et al., 1996; Andreasen et al., 1998], or those with cerebellar infarct [Schmahmann and Sherman, 1998].

Given the complex nature of EF, as well as its importance in the completion of activities of daily living, maintaining this domain is critically important. Here, the suggestion that the cerebellum engages with the PFC in a manner that parallels the gradient of abstraction in the PFC means that we now have additional areas of investigation for understanding EF, or its breakdown in the case of disease or infarct. Further, this provides additional targets of intervention and remediation to improve these skills in impacted populations. Future work targeting the relative contributions of the cerebellum, and ideally its lobular subregions, taking advantage of non-invasive brain stimulation stand to provide important insights into the necessity of this region for EF, and its underlying domains of function.

## Acknowledgements

Data were provided by the Human Connectome Project, WU-Minn Consortium (Principal Investigators: David Van Essen and Kamil Ugurbil; 1U54MH091657) funded by the 16 NIH Institutes and Centers that support the NIH Blue-print for Neuroscience Research; and by the McDonnell Center for Systems Neuroscience at Washington University. The authors acknowledge the Texas A&M University Brazos HPC cluster that contributed to the research reported here. Results are accessible via the Brain Analysis Library of Spatial Maps and Atlases (https://balsa.wustl.edu/study/show/k53g). Analysis scripts and other results are available here: https://osf.io/3vka4/.

