## Supplemental Figures & Tables for "Dissociable Cerebellar-Prefrontal Networks Underlying Executive Function: Evidence from the Human Connectome Project"

### Supplemental Figures and Tables

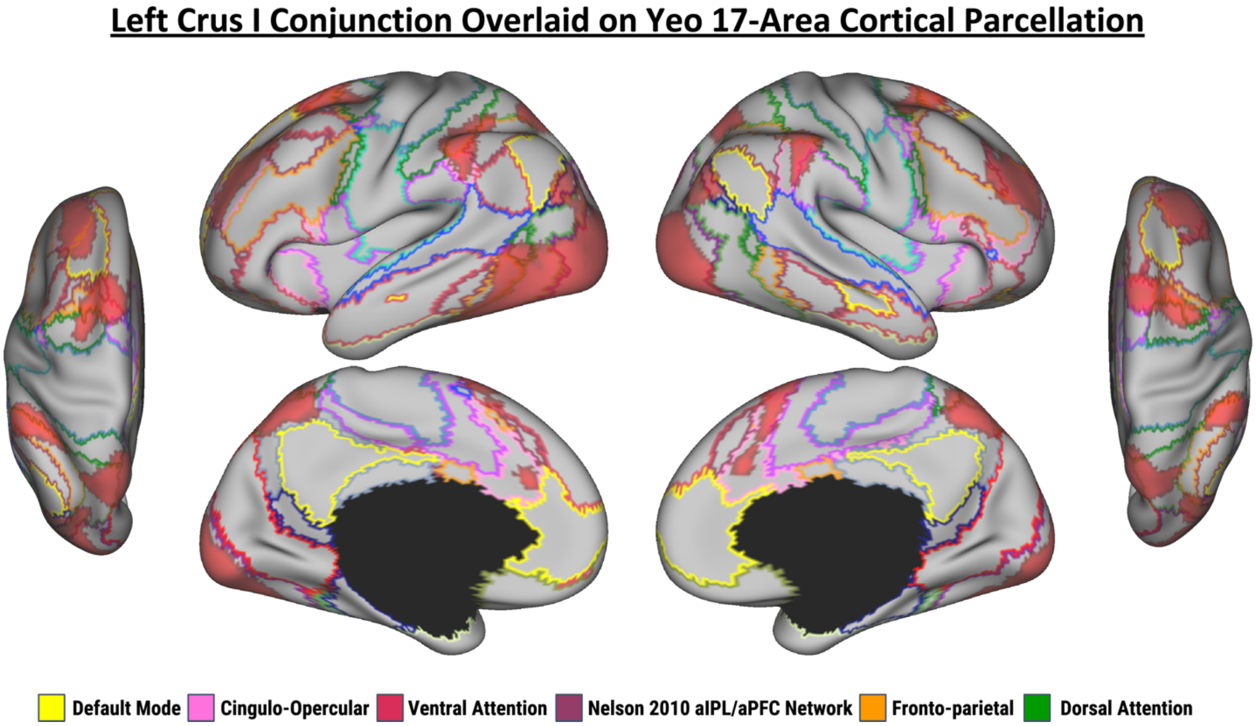

Figure S1. Resting state functional connectivity results from the conjunction of Left Crus I contrasts. Connectivity is overlaid on the outline of Yeo et al. [2011] 17-Area Cortical Parcellation.

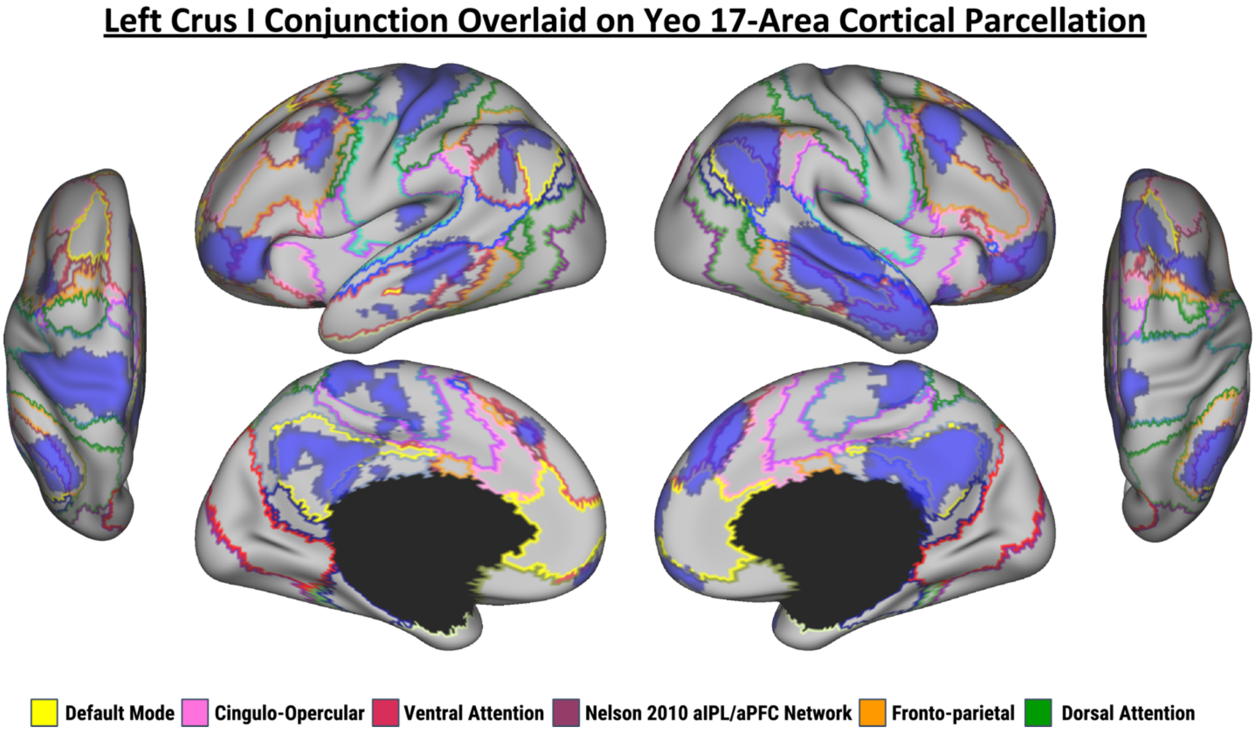

Figure S2. Resting state functional connectivity results from the conjunction of Crus II contrasts. Connectivity is overlaid on the outline of Yeo et al. [2011] 17-Area Cortical Parcellation.

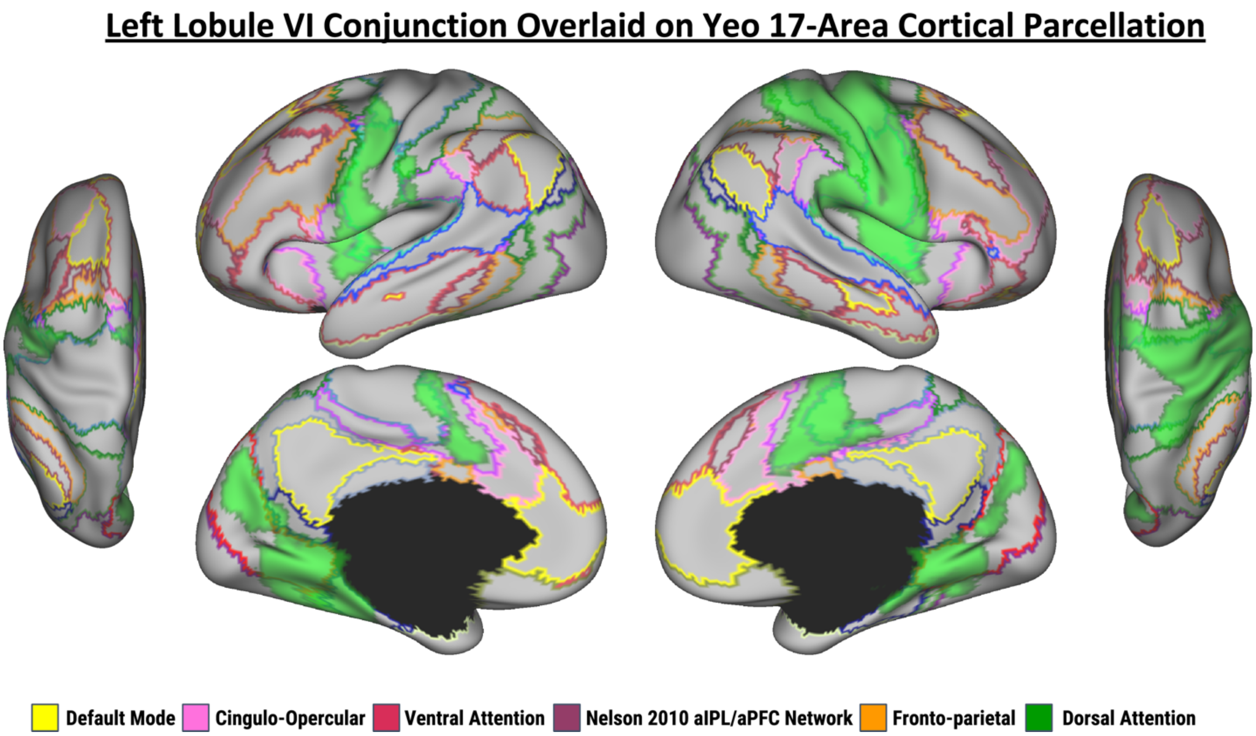

Figure S3. Resting state functional connectivity results from the conjunction of Lobule VI contrasts. Connectivity is overlaid on the outline of the Yeo et al. [2011] 17-Area Cortical Parcellation.

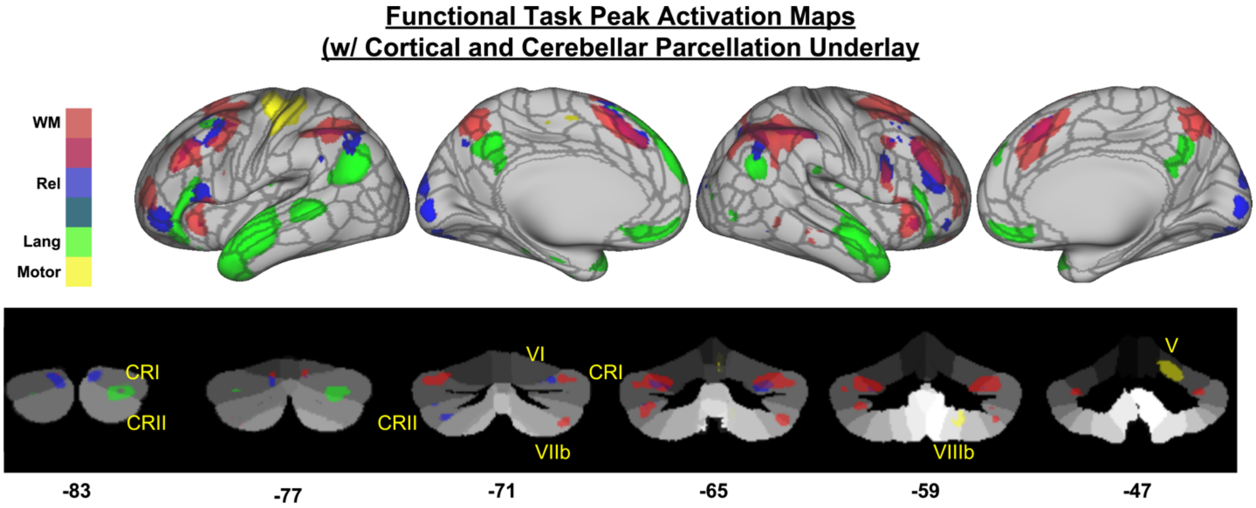

Figure S4. Peak activations from the four functional tasks: working memory (red), relational processing (blue), language (green), and motor (yellow) overlaid on the inflated cortical surface (top row in each section) and the cerebellum (bottom row in each section). Cluster-corrected activation maps were further thresholded at the 95 percentile to isolate peaks of activation, and were binarized for display. The working memory activation was thresholded at T>10.98, relational processing was thresholded at T>9.34, language was thresholded at T>15.5, and motor was thresholded at T>13.45. Section A shows the activation maps on the template surface/volume, while section B shows the activation maps with the Glasser et al. (2016) 360 region cortical parcellation shown on the cortical surfaces and the Diedrichsen Cerebellar SUIT atlas.

Supplemental Table 1. Atlas query of conjunction of Left Crus I - Crus II connectivity results. For large clusters, sub-region labels are listed below. COG: Center of Gravity.

| ClusterIndex | Voxels | Max T-Value | COG X (mm) | COG Y (mm) | COG Z (mm) | COG Label |
|---|---|---|---|---|---|---|
| 9 | 23218 | 37.1 | -7.51 | -67.6 | -3.76 | 73% Lingual Gyrus |
| 8 | 4187 | 12.3 | 26 | 10.7 | 41.6 | 28% Anterior PMd (Premotor dorsal) |
| 7 | 1531 | 14.3 | 35.1 | 44.6 | 26.7 | 50% Area 9/46D |
| 6 | 1166 | 11.6 | -31.3 | 44.9 | 28.7 | 63% Area 9/46D |
| 5 | 959 | 9.19 | -21.8 | 5.6 | 61.7 | 45% Anterior PMd (Premotor dorsal) |
| 4 | 104 | 7.1 | 10.1 | -77.3 | -42.7 | 71% Right Crus II |
| 3 | 72 | 6.11 | -45.4 | 16.3 | -1.85 | 43% Frontal Operculum Cortex |
| 2 | 70 | 6.1 | -46.4 | 2.95 | 33.7 | 50% Precentral Gyrus |
| 1 | 41 | 5.97 | -41.3 | -1.76 | 49 | 61% Area 9/46D |

-----------------------------------------

Structures to which each cluster belongs to:

| Cluster #9 | Average Atlas Probability Within Cluster |
|---|---|
| Lateral Occipital Cortex, superior division | 10.0 |
| Lateral Occipital Cortex, inferior division | 8.7 |
| Occipital Pole | 6.6 |

| Cluster #8 | Average Atlas Probability Within Cluster |
|---|---|
| Supplementary Motor Area | 6.2 |
| Anterior PMd (Premotor dorsal) | 8.3 |
| Precentral Gyrus | 9.8 |
| Paracingulate Gyrus | 9.6 |
| Cingulate Gyrus, anterior division | 7.8 |

| Cluster #7 | Average Atlas Probability Within Cluster |
|---|---|
| Area 9/46D | 21.5 |
| Area 9/46V | 12.0 |
| Area 46 | 17.7 |

| Cluster #6 | Average Atlas Probability Within Cluster |
|---|---|
| Area 9/46D | 33.1 |
| Area 46 | 21.2 |

| Cluster #5 | Average Atlas Probability Within Cluster |
|---|---|
| Superior Frontal Gyrus | 29.9 |
| Middle Frontal Gyrus | 8.5 |

Supplemental Table 2. Atlas query of conjunction of Right Crus I - Crus II connectivity results. For large clusters, sub-region labels are listed below. COG: Center of Gravity.

| ClusterIndex | Voxels | MAX | COG X (mm) | COG Y (mm) | COG Z (mm) | COG Label |
|---|---|---|---|---|---|---|
| 16 | 3872 | 39.4 | 38.2 | -65.8 | -27.3 | 99% Right Crus I |
| 15 | 3831 | 15.8 | -1.77 | 14.8 | 57.3 | 30% Cluster1 (SMA) |
| 14 | 1336 | 13.7 | -30 | 46.7 | 26.6 | 54% Cluster5 (Area 9/46D) |
| 13 | 1205 | 14.9 | -46.6 | 17.8 | 1.38 | 100% 44v |
| 12 | 663 | 9.18 | 31.4 | 50.9 | 26.4 | 67% Cluster7 (Area 46) |
| 11 | 407 | 10.9 | -38.4 | -62.8 | -27.6 | 100% Left Crus I |
| 10 | 269 | 8.24 | 46.2 | 19.9 | -2.66 | 44% Frontal Operculum Cortex |
| 9 | 252 | 6.85 | -56.5 | -46.8 | 32.5 | 48% Supramarginal Gyrus |
| 8 | 209 | 8.81 | -28.8 | 57.3 | -12.8 | 87% FPl |
| 7 | 164 | 6.67 | -49.5 | 1.98 | 48.4 | 84% 6v |
| 6 | 98 | 6.6 | -46.3 | -55.2 | -44.5 | 59% Left Crus II |
| 5 | 97 | 6.81 | -31.1 | -1.2 | 62.1 | 41% Cluster8 (Area 8A) |
| 4 | 56 | 5.93 | 59 | -50.4 | 43.1 | 63% Angular Gyrus |
| 3 | 8 | 5.33 | 48.5 | -40.5 | 37.5 | 33% Supramarginal Gyrus |
| 2 | 5 | 5.72 | 3.19 | -96.8 | 19.2 | 42% Occipital Pole |
| 1 | 1 | 5.01 | 4 | -98 | -10 | 19% Occipital Pole |

-----------------------------------------

Structures to which each cluster belongs to:

| Cluster #16 | Average Atlas Probability Within Cluster |
|---|---|
| Right VI | 10.9 |
| Right Crus I | 34.7 |
| Lateral Occipital Cortex, inferior division | 6.8 |
| Occipital Fusiform Gyrus | 5.7 |

| Cluster #15 | Average Atlas Probability Within Cluster |
|---|---|
| Cluster1 (SMA) | 11.5 |
| Paracingulate Gyrus | 15.6 |
| Cingulate Gyrus, anterior division | 6.5 |

| Cluster #14 | Average Atlas Probability Within Cluster |
|---|---|
| Cluster5 (Area 9/46D) | 28.5 |
| Cluster6 (Area 9/46V) | 8.9 |
| Cluster7 (Area 46) | 25.6 |

| Cluster #13 | Average Atlas Probability Within Cluster |
|---|---|
| 6r | 10.0 |
| 44v | 12.2 |
| Frontal Orbital Cortex | 11.6 |
| Frontal Operculum Cortex | 11.9 |

| Cluster #12 | Average Atlas Probability Within Cluster |
|---|---|
| Area 46 | 17.7 |
| Cluster5 (Area 9/46D) | 19.8 |
| Cluster7 (Area 46) | 26.1 |

| Cluster #11 | Average Atlas Probability Within Cluster |
|---|---|
| Left VI | 21.0 |
| Left Crus I | 69.6 |

| Cluster #10 | Average Atlas Probability Within Cluster |
|---|---|
| 6r | 7.5 |
| 44v | 18.0 |
| Frontal Orbital Cortex | 21.2 |
| Frontal Operculum Cortex | 16.3 |

| Cluster #9 | Average Atlas Probability Within Cluster |
|---|---|
| Supramarginal Gyrus, posterior division | 38.6 |
| Angular Gyrus | 14.4 |

| Cluster #6 | Average Atlas Probability Within Cluster |
|---|---|
| Left Crus I | 29.2 |
| Left Crus II | 47.5 |

| Cluster #5 | Average Atlas Probability Within Cluster |
|---|---|
| Cluster8 (Area 8A) | 25.7 |
| Cluster9 (Ant PMd) | 31.0 |

Supplemental Table 3. Atlas query of conjunction of Left Crus I - Lobule VI connectivity results. For large clusters, sub-region labels are listed below. COG: Center of Gravity.

| ClusterIndex | Voxels | Max T-Value | COG X (mm) | COG Y (mm) | COG Z (mm) | COG Label |
|---|---|---|---|---|---|---|
| 18 | 8582 | 35.9 | -28.2 | -75 | -23.8 | 81% Left Crus I |
| 17 | 1143 | 9.24 | 1.78 | -67.6 | 59.6 | 20% Precuneous Cortex |
| 16 | 430 | 8.08 | 21.4 | 14.3 | 63.8 | 41% Anterior PMd (Premotor dorsal) |
| 15 | 250 | 7.1 | 51.8 | -40.9 | 46.7 | 53% Supramarginal Gyrus, posterior division |
| 14 | 221 | 7.3 | 40.4 | -77.5 | 32.2 | 76% Lateral Occipital Cortex, superior division |
| 13 | 202 | 6.74 | 30.4 | 52.9 | 29.9 | 41% Area 46 |
| 12 | 192 | 6.35 | -19.5 | 13.1 | 64.7 | 41% Supplemental Motor Area |
| 11 | 184 | 6.99 | 40.6 | -45.7 | -42.7 | 50% Right Crus II |
| 10 | 57 | 6.08 | -58.2 | -41.3 | 46.7 | 43% Supramarginal Gyrus, posterior division |
| 9 | 44 | 5.82 | -28.7 | 49.2 | 35.7 | 44% Area 9/46D |
| 8 | 36 | 5.78 | -35.1 | -86.5 | 32.3 | 63% Lateral Occipital Cortex |

------------------------------------------

Structures to which each cluster belongs to:

| Cluster #18 | Average Atlas Probability Within Cluster |
|---|---|
| Left Crus I | 19.0 |
| Lateral Occipital Cortex, inferior division | 8.0 |
| Occipital Pole | 13.1 |

| Cluster #17 | Average Atlas Probability Within Cluster |
|---|---|
| Lateral Occipital Cortex, superior division | 33.1 |
| Precuneous Cortex | 19.4 |

| Cluster #16 | Average Atlas Probability Within Cluster |
|---|---|
| Area 8A | 13.1 |
| Anterior PMd (Premotor dorsal) | 13.9 |

| Cluster #15 | Average Atlas Probability Within Cluster |
|---|---|
| Supramarginal Gyrus, posterior division | 42.8 |
| Angular Gyrus | 11.7 |

| Cluster #12 | Average Atlas Probability Within Cluster |
|---|---|
| Supplemental Motor Area | 18.3 |
| Pre-Supplemental Motor Area | 11.3 |
| Anterior PMd (Premotor dorsal) | 20.3 |

| Cluster #11 | Average Atlas Probability Within Cluster |
|---|---|
| Right Crus I | 23.1 |
| Right Crus II | 32.5 |
| Right VIIb | 7.2 |

| Cluster #10 | Average Atlas Probability Within Cluster |
|---|---|
| Supramarginal Gyrus, anterior division | 28.2 |
| Supramarginal Gyrus, posterior division | 33.8 |

| Cluster #8 | Average Atlas Probability Within Cluster |
|---|---|
| Lateral Occipital Cortex, superior division | 51.5 |
| Occipital Pole | 11.4 |

| Cluster #7 | Average Atlas Probability Within Cluster |
|---|---|
| Superior Parietal Lobule | 16.7 |
| Supramarginal Gyrus, anterior division | 7.7 |
| Supramarginal Gyrus, posterior division | 36.8 |

| Supplemental Table 4. Atlas query of conjunction of Right Crus I - Lobule VI connectivity results. For large clusters, sub-region labels are listed below. COG: Center of Gravity. | | | | | | |
|---|---|---|---|---|---|---|
| ClusterIndex | Voxels | MAX | COG X (mm) | COG Y (mm) | COG Z (mm) | COG Label |
| 16 | 4150 | 37.8 | 36 | -72.1 | -28.6 | 98% Right Crus I |
| 15 | 2283 | 12.5 | -1.35 | 21.7 | 60.1 | 28% Pre-Supplementary Motor Area |
| 14 | 816 | 9.08 | -46.1 | 22.8 | -1.9 | 51% Frontal Operculum Cortex |
| 13 | 571 | 8.51 | -54.6 | -52.5 | 39 | 39% Angular Gyrus |
| 12 | 265 | 7.46 | -33.5 | -78.4 | -29.2 | 95% Left Crus I |
| 11 | 209 | 7.16 | 40.9 | 21.5 | -5.82 | 49% Frontal Orbital Cortex |
| 10 | 202 | 6.98 | -30.8 | 59.9 | -8.15 | 96% FPl (Frontal Pole lateral - ventral) |
| 9 | 201 | 6.55 | 55.3 | -49 | 43.1 | 67% Angular Gyrus |
| 8 | 191 | 6.79 | -24.6 | 54.2 | 29.6 | 41% Area 9/46D |
| 7 | 142 | 9.19 | 43.9 | -59.4 | -53 | 57% Right VIIb |
| 6 | 114 | 6.86 | -45.2 | 15.3 | 47.9 | 32% Area 8A |
| 5 | 95 | 7.08 | 3.35 | -38.1 | 75.4 | 38% Postcentral Gyrus |
| 4 | 67 | 6.3 | 51.6 | 36.3 | -9.06 | 54% Frontal Pole |
| 3 | 51 | 7.38 | -29.9 | -55 | -31.2 | 97% Left VI |
| ----------------------------------------- |  |  |  |  |  |  |
| Structures to which each cluster belongs to: | | | | | | |
| Cluster #16 | | | Average Atlas Probability Within Cluster | | | |
| Right Crus I | | | 43.1 | | | |
| Right Crus II | | | 10.1 | | | |
| Cluster #15 | | | Average Atlas Probability Within Cluster | | | |
| Supplementary Motor Area | | | 7.3 | | | |
| Pre-Supplementary Motor Area | | | 11.2 | | | |
| Paracingulate Gyrus | | | 5.1 | | | |
| Cluster #14 | | | Average Atlas Probability Within Cluster | | | |
| 44v | | | 13.2 | | | |
| 45A | | | 5.3 | | | |
| Frontal Orbital Cortex | | | 18.3 | | | |
| Frontal Operculum Cortex | | | 8.5 | | | |
| Cluster #13 | | | Average Atlas Probability Within Cluster | | | |
| Supramarginal Gyrus, posterior division | | | 27.3 | | | |
| Angular Gyrus | | | 27.6 | | | |
| Lateral Occipital Cortex, superior division | | | 8.1 | | | |
| Cluster #12 | | | Average Atlas Probability Within Cluster | | | |
| Left Crus I | | | 85.1 | | | |
| Left Crus II | | | 8.8 | | | |
| Cluster #11 | | | Average Atlas Probability Within Cluster | | | |
| Insular Cortex | | | 14.1 | | | |
| Area 44v | | | 8.3 | | | |
| Fop (Frontal Opercular) | | | 9.2 | | | |
| Frontal Orbital Cortex | | | 36.6 | | | |
| Frontal Operculum Cortex | | | 9.8 | | | |
| Cluster #9 | | | Average Atlas Probability Within Cluster | | | |
| Supramarginal Gyrus, posterior division | | | 18.0 | | | |
| Angular Gyrus | | | 42.8 | | | |
| Cluster #7 | | | Average Atlas Probability Within Cluster | | | |
| Right Crus II | | | 29.5 | | | |
| Right VIIb | | | 44.9 | | | |
| Cluster #5 | | | Average Atlas Probability Within Cluster | | | |
| Precentral Gyrus | | | 7.6 | | | |
| Postcentral Gyrus | | | 27.7 | | | |
| Cluster #4 | | | Average Atlas Probability Within Cluster | | | |
| Area 45A | | | 12.2 | | | |
| Area 47 | | | 31.6 | | | |
| IFS (Inferior Frontal Sulcus) | | | 14.1 | | | |

Supplemental Table 5. Atlas query of conjunction of Left Crus II - Crus I connectivity results. For large clusters, sub-region labels are listed below. COG: Center of Gravity.

| ClusterIndex | Voxels | MAX | COG X (mm) | COG Y (mm) | COG Z (mm) | COG Label |
|---|---|---|---|---|---|---|
| 16 | 5433 | 9.63 | -39.9 | -19.4 | 39.6 | 60% Postcentral Gyrus |
| 15 | 4348 | 9.43 | 45.3 | -16.5 | 37.6 | 47% Postcentral Gyrus |
| 14 | 2665 | 38.8 | -25.8 | -77.3 | -37.7 | 94% Left Crus II |
| 13 | 2074 | 10.4 | 62.8 | -19.7 | -12.2 | 51% Middle Temporal Gyrus, posterior division |
| 12 | 1834 | 10.7 | 3.1 | -57.4 | 34 | 86% Precuneous Cortex |
| 11 | 1279 | 8.08 | -12.2 | 59.4 | -5.07 | Area 10 (Dorsal) |
| 10 | 1197 | 14.8 | 49.8 | -59.9 | 37.1 | 48% Lateral Occipital Cortex, superior division |
| 9 | 1196 | 12.6 | 28.4 | -79 | -33.5 | 91% Right Crus I |
| 8 | 1002 | 9.77 | 7.98 | 42.9 | 45.5 | 43% Area 9 |
| 7 | 511 | 9.22 | -41.9 | -66.4 | 44.8 | 66% Lateral Occipital Cortex, superior division |
| 6 | 478 | 11.6 | -16 | -57.8 | -20.6 | 87% Left VI |
| 5 | 445 | 7.74 | 46 | 43.3 | -11.4 | 58% Area 47 |
| 4 | 311 | 10.3 | -5.26 | -60.1 | -47.5 | 85% Left IX |
| 3 | 253 | 8.17 | 42.3 | 18.1 | 46 | 34% Area 8A |
| 2 | 239 | 6.72 | -41.4 | 17.1 | 45.4 | 34% Area 8A |
| 1 | 176 | 6.51 | -62.8 | -30.4 | -6.61 | 59% Middle Temporal Gyrus, posterior division |

-----------------------------------------

Structures to which each cluster belongs to:

| Cluster #16 | Average Atlas Probability Within Cluster |
|---|---|
| Precentral Gyrus | 16.3 |
| Postcentral Gyrus | 23.2 |
| Central Opercular Cortex | 7.0 |

| Cluster #15 | Average Atlas Probability Within Cluster |
|---|---|
| Precentral Gyrus | 16.2 |
| Postcentral Gyrus | 21.0 |
| Central Opercular Cortex | 8.7 |
| Parietal Operculum Cortex | 4.1 |

| Cluster #14 | Average Atlas Probability Within Cluster |
|---|---|
| Left Crus I | 25.4 |
| Left Crus II | 47.4 |

| Cluster #12 | Average Atlas Probability Within Cluster |
|---|---|
| Cingulate Gyrus, posterior division | 24.6 |
| Precuneous Cortex | 42.2 |
| Cuneal Cortex | 5.5 |

| Cluster #11 | Average Atlas Probability Within Cluster |
|---|---|
| Frontal Pole Medial (ventral) | 18.7 |
| Area 10 (Dorsal) | 7.9 |
| Area 46 | 7.4 |

| Cluster #10 | Average Atlas Probability Within Cluster |
|---|---|
| Angular Gyrus | 26.9 |
| Lateral Occipital Cortex, superior division | 32.9 |

| Cluster #9 | Average Atlas Probability Within Cluster |
|---|---|
| Right Crus I | 49.8 |
| Right Crus II | 33.7 |

| Cluster #8 | Average Atlas Probability Within Cluster |
|---|---|
| Area 9 | 13.2 |
| Area 8B | 12.0 |
| Pre-Supplementary Motor Area | 8.4 |
| Paracingulate Gyrus | 3.0 |

| Cluster #7 | Average Atlas Probability Within Cluster |
|---|---|
| Angular Gyrus | 8.4 |
| Lateral Occipital Cortex, superior division | 53.1 |

| Cluster #6 | Average Atlas Probability Within Cluster |
|---|---|
| Left V | 26.9 |
| Left VI | 65.4 |

| Cluster #3 | Average Atlas Probability Within Cluster |
|---|---|
| Area 9/46V | 17.2 |
| Area 8A | 13.0 |

| Cluster #2 | Average Atlas Probability Within Cluster |
|---|---|
| Area 9/46V | 20.6 |
| Area 8A | 17.0 |

Supplemental Table 6. Atlas query of conjunction of Right Crus II - Crus I connectivity results. For large clusters, sub-region labels are listed below. COG: Center of Gravity.

| ClusterIndex | Voxels | Max T-Value | COG X (mm) | COG Y (mm) | COG Z (mm) | COG Label |
|---|---|---|---|---|---|---|
| 17 | 4749 | 12.5 | -1.63 | -54 | 28.2 | 64% Cingulate Gyrus |
| 16 | 4585 | 38.6 | 20.4 | -72 | -40.1 | 21% Right Crus II |
| 15 | 2417 | 12.8 | -2.27 | 56.6 | -3.66 | 44% Frontal Pole |
| 14 | 2106 | 11.8 | -39.6 | -72.1 | 36.3 | 60% Lateral Occipital Cortex, superior division |
| 13 | 1643 | 8.85 | -54 | -14.2 | -2.48 | 15% Superior Temporal Gyrus, posterior division |
| 12 | 1060 | 10.3 | -59.7 | -50.6 | -9.44 | 51% Middle Temporal Gyrus, temporooccipital part |
| 11 | 1012 | 9.1 | 60.9 | -20.9 | 17.3 | 28% Parietal Operculum Cortex |
| 10 | 695 | 8.73 | -22.8 | 27.3 | 47.7 | 58% Cluster10 (area 8B) |
| 9 | 503 | 6.76 | 48.2 | -64.8 | 31.2 | 68% Lateral Occipital Cortex, superior division |
| 8 | 351 | 6.99 | -43.3 | 26 | 22 | 43% Cluster6 (Area 9/46V) |
| 7 | 335 | 11.2 | -33.1 | 34.4 | -12.8 | 96% Fop |
| 6 | 308 | 7.22 | 43.3 | 26.2 | 20.5 | 42% Cluster6 (Area 9/46V) |
| 5 | 251 | 7.71 | -28.7 | -35 | -17.7 | 51% Parahippocampal Gyrus, posterior division |
| 4 | 220 | 8.12 | 58.6 | -5.15 | -14.9 | 27% Middle Temporal Gyrus, anterior division |
| 3 | 177 | 9.15 | -22.4 | -19.4 | -15.9 | 3% Parahippocampal Gyrus |
| 2 | 104 | 9.53 | -11.9 | -71.2 | -25 | 94% Left VI |
| 1 | 90 | 7.1 | 23.6 | -20 | -16.4 | 90% Right Hippocampus |

-----------------------------------------

Structures to which each cluster belongs to:

| Cluster #17 | Average Atlas Probability Within Cluster |
|---|---|
| Cingulate Gyrus, posterior division | 21.4 |
| Precuneous Cortex | 37.2 |
| Cuneal Cortex | 2.7 |
| Supracalcarine Cortex | 2.4 |

| Cluster #15 | Average Atlas Probability Within Cluster |
|---|---|
| FPm | 9.1 |
| Cluster4 (Area 10) | 20.0 |
| Frontal Medial Cortex | 17.4 |
| Paracingulate Gyrus | 11.1 |

| Cluster #14 | Average Atlas Probability Within Cluster |
|---|---|
| Lateral Occipital Cortex, superior division | 58.1 |

| Cluster #13 | Average Atlas Probability Within Cluster |
|---|---|
| Superior Temporal Gyrus, anterior division | 6.8 |
| Superior Temporal Gyrus, posterior division | 8.5 |
| Middle Temporal Gyrus, anterior division | 8.3 |
| Middle Temporal Gyrus, posterior division | 7.0 |
| Heschl's Gyrus (includes H1 and H2) | 7.3 |
| Planum Temporale | 10.8 |

| Cluster #12 | Average Atlas Probability Within Cluster |
|---|---|
| Middle Temporal Gyrus, posterior division | 8.1 |
| Middle Temporal Gyrus, temporooccipital part | 26.1 |
| Inferior Temporal Gyrus, temporooccipital part | 16.3 |

| Cluster #11 | Average Atlas Probability Within Cluster |
|---|---|
| Superior Temporal Gyrus, posterior division | 7.4 |
| Postcentral Gyrus | 9.9 |
| Supramarginal Gyrus, anterior division | 8.2 |
| Central Opercular Cortex | 11.3 |
| Parietal Operculum Cortex | 13.5 |
| Planum Temporale | 13.4 |

| Cluster #10 | Average Atlas Probability Within Cluster |
|---|---|
| Cluster2 (PreSMA) | 12.6 |
| Cluster5 (Area 9/46D) | 6.5 |
| Cluster8 (Area 8A) | 5.4 |
| Cluster10 (area 8B) | 36.6 |

| Cluster #8 | Average Atlas Probability Within Cluster |
|---|---|
| IFJ | 14.2 |
| 44d | 21.6 |
| IFS | 13.7 |
| Cluster6 (Area 9/46V) | 26.7 |

| Cluster #7 | Average Atlas Probability Within Cluster |
|---|---|
| Fop | 24.0 |
| Area 47 | 5.9 |
| Frontal Orbital Cortex | 37.4 |

| Cluster #6 | Average Atlas Probability Within Cluster |
|---|---|
| 44d | 22.3 |
| IFS | 18.7 |
| Cluster6 (Area 9/46V) | 25.1 |

| Cluster #5 | Average Atlas Probability Within Cluster |
|---|---|
| Parahippocampal Gyrus, posterior division | 32.5 |
| Temporal Fusiform Cortex, posterior division | 32.9 |

| Cluster #4 | Average Atlas Probability Within Cluster |
|---|---|
| Superior Temporal Gyrus, anterior division | 18.3 |
| Superior Temporal Gyrus, posterior division | 12.2 |
| Middle Temporal Gyrus, anterior division | 23.1 |
| Middle Temporal Gyrus, posterior division | 15.9 |

Supplemental Table 7. Atlas query of conjunction of Left Crus II - Lobule VI connectivity results. For large clusters, sub-region labels are listed below. COG: Center of Gravity.

| ClusterIndex | Voxels | Max T-Value | COG X (mm) | COG Y (mm) | COG Z (mm) | COG Label |
|---|---|---|---|---|---|---|
| 18 | 3869 | 44.7 | -27.7 | -76.5 | -36.9 | 73% Left Crus I |
| 17 | 1811 | 10.9 | 62.7 | -24.8 | -11.3 | 54% Middle Temporal Gyrus, posterior division |
| 16 | 1778 | 10.7 | 14.8 | 38.3 | 46.9 | 54% Area 8B |
| 15 | 1760 | 13.8 | 49.8 | -58 | 38 | 36% Angular Gyrus |
| 14 | 1450 | 10 | 4.69 | -54 | 37.1 | 76% Precuneous Cortex |
| 13 | 1013 | 10.6 | 28.4 | -79 | -32.5 | 95% Right Crus I |
| 12 | 819 | 7.51 | -34.2 | -27.9 | 63.5 | 38% Postcentral Gyrus |
| 11 | 360 | 6.97 | 47 | 45.8 | -8.86 | 55% Area 47 |
| 10 | 231 | 9.7 | -3.86 | -56 | -48 | 99% Left IX |
| 9 | 197 | 6.47 | -1.5 | -39.4 | 64.9 | 42% Postcentral Gyrus |
| 8 | 173 | 6.35 | -64.1 | -34.3 | -6.16 | 71% Middle Temporal Gyrus, posterior division |
| 7 | 157 | 6.82 | -41.6 | -65.9 | 50.8 | 72% Lateral Occipital Cortex, superior division |
| 6 | 109 | 6.54 | 13 | 69 | 14.5 | 61% Frontal Pole |
| 5 | 97 | 8.61 | 24.2 | -49.7 | -26.7 | 82% Right VI |
| 4 | 96 | 7.2 | 7.38 | 11.1 | 11.4 | 54% Right Caudate |
| 3 | 93 | 6.3 | -35.4 | 58 | -2.13 | 81% FPl (Frontal Pole Lateral) |
| 2 | 77 | 6.83 | -41.6 | 16.1 | 39.8 | 44% Cluster6 Area 9/46V |

------------------------------------------

Structures to which each cluster belongs to:

| Cluster #18 | Average Atlas Probability Within Cluster |
|---|---|
| Left Crus I | 32.0326 |
| Left Crus II | 36.6875 |
| Cluster #17 | Average Atlas Probability Within Cluster |
| Middle Temporal Gyrus, posterior division | 31.312 |
| Middle Temporal Gyrus, temporooccipital part | 7.9536 |
| Cluster #16 | Average Atlas Probability Within Cluster |
| Pre-Supplementary Motor Area | 8.2499 |
| Area 9 | 7.887 |
| Area 8B | 9.9026 |
| Cluster #15 | Average Atlas Probability Within Cluster |
| Angular Gyrus | 30.0347 |
| Lateral Occipital Cortex, superior division | 26.2631 |
| Cluster #14 | Average Atlas Probability Within Cluster |
| Cingulate Gyrus, posterior division | 25.9097 |
| Precuneous Cortex | 44.1317 |
| Cluster #13 | Average Atlas Probability Within Cluster |
| Right Crus I | 57.7848 |
| Right Crus II | 29.4669 |
| Cluster #12 | Average Atlas Probability Within Cluster |
| Precentral Gyrus | 15.265 |
| Postcentral Gyrus | 38.4579 |
| Cluster #11 | Average Atlas Probability Within Cluster |
| Area 47 | 18 |
| IFS (Inferior Frontal Sulcus) | 7.1472 |
| Cluster #10 | Average Atlas Probability Within Cluster |
| Left IX | 69.1602 |
| Right IX | 12.1732 |
| Cluster #9 | Average Atlas Probability Within Cluster |
| Precentral Gyrus | 12.4416 |
| Postcentral Gyrus | 34.3452 |

Supplemental Table 8. Atlas query of conjunction of Right Crus II - Lobule VI connectivity results. For large clusters, sub-region labels are listed below. COG: Center of Gravity.

| ClusterIndex | Voxels | MAX | COG X (mm) | COG Y (mm) | COG Z (mm) | COG Label |
|---|---|---|---|---|---|---|
| 12 | 3802 | 41.7 | 29.3 | -75.1 | -38.2 | 83% Right Crus II |
| 11 | 1773 | 13.9 | -41.9 | -68.5 | 39 | 57% Lateral Occipital Cortex, superior division |
| 10 | 1594 | 10.6 | -20.6 | 31.5 | 49.4 | 55% Area 8B |
| 9 | 1422 | 9.99 | -3.75 | -46.6 | 32.5 | 78% Cingulate Gyrus, posterior division |
| 8 | 980 | 11.1 | -61.2 | -45.8 | -8.44 | 50% Middle Temporal Gyrus, temporooccipital part |
| 7 | 841 | 8.62 | -3.5 | 62.3 | 1.74 | 64% Area 10 (dorsal) |
| 6 | 599 | 8.01 | -58.3 | -15.1 | -17.7 | 38% Middle Temporal Gyrus, posterior division |
| 5 | 378 | 7.17 | -43.3 | 42.8 | -9.41 | 60% Area 47 |
| 4 | 327 | 10.8 | 3.81 | -53.5 | -48.1 | 95% Right IX |
| 3 | 243 | 7.6 | -20.3 | -87 | -34.5 | 99% Left Crus II |
| 2 | 191 | 8.51 | -42.4 | -75 | -34.3 | 97% Left Crus I |

-----------------------------------------

Structures to which each cluster belongs to:

| Cluster #12 | Average Atlas Probability Within Cluster |
|---|---|
| Right Crus I | 29.8 |
| Right Crus II | 38.6 |

| Cluster #11 | Average Atlas Probability Within Cluster |
|---|---|
| Angular Gyrus | 9.1 |
| Lateral Occipital Cortex, superior division | 52.3 |

| Cluster #10 | Average Atlas Probability Within Cluster |
|---|---|
| Pre-Supplementary Motor Area | 10.8 |
| Area 9 | 5.4 |
| Area 8B | 20.2 |

| Cluster #9 | Average Atlas Probability Within Cluster |
|---|---|
| Cingulate Gyrus, posterior division | 41.7 |
| Precuneous Cortex | 29.0 |

| Cluster #8 | Average Atlas Probability Within Cluster |
|---|---|
| Middle Temporal Gyrus, posterior division | 19.8 |
| Middle Temporal Gyrus, temporooccipital part | 23.4 |
| Inferior Temporal Gyrus, posterior division | 3.7 |
| Inferior Temporal Gyrus, temporooccipital part | 8.6 |

| Cluster #7 | Average Atlas Probability Within Cluster |
|---|---|
| FPm (Frontal Pole medial - ventral) | 19.9 |
| Area 10 (dorsal) | 24.8 |
| Frontal Medial Cortex | 10.5 |
| Paracingulate Gyrus | 4.5 |

| Cluster #6 | Average Atlas Probability Within Cluster |
|---|---|
| Superior Temporal Gyrus, posterior division | 6.6 |
| Middle Temporal Gyrus, anterior division | 13.5 |
| Middle Temporal Gyrus, posterior division | 31.7 |
| Inferior Temporal Gyrus, posterior division | 4.0 |

| Cluster #5 | Average Atlas Probability Within Cluster |
|---|---|
| Fop (Frontal operculus) | 6.7 |
| Area 47 | 21.7 |
| IFS (Inferior Frontal Sulcus) | 14.3 |
| FPl (Frontal Pole lateral - ventral) | 5.1 |
| Area 46 | 4.6 |

| Cluster #4 | Average Atlas Probability Within Cluster |
|---|---|
| Left IX | 15.9 |
| Right IX | 65.1 |

| Cluster #3 | Average Atlas Probability Within Cluster |
|---|---|
| Left Crus I | 29.5 |
| Left Crus II | 52.8 |

Supplemental Table 9. Atlas query of conjunction of Left Lobule VI - Crus I connectivity results. For large clusters, sub-region labels are listed below. COG: Center of Gravity.

| ClusterIndex | Voxels | MAX | COG X (mm) | COG Y (mm) | COG Z (mm) | COG Label |
|---|---|---|---|---|---|---|
| 4 | 11809 | 15.8 | 40.8 | -13.2 | 37.4 | 34% Precentral Gyrus |
| 3 | 6997 | 50.7 | -13.3 | -59.5 | -20 | 91% Left VI |
| 2 | 4141 | 12.8 | -52.3 | -9.76 | 22.6 | 21% Postcentral Gyrus |
| 1 | 132 | 6.73 | -21.5 | -31.5 | 63 | 45% Postcentral Gyrus |

------------------------------------------

Structures to which each cluster belongs to:

| Cluster #4 | Average Atlas Probability Within Cluster |
|---|---|
| Precentral Gyrus | 12.8 |
| Postcentral Gyrus | 12.5 |
| Juxtapositional Lobule Cortex (formerly Supplem | 5.0 |
| Central Opercular Cortex | 5.3 |

| Cluster #3 | Average Atlas Probability Within Cluster |
|---|---|
| Left V | 8.5 |
| Left VI | 14.7 |
| Precuneous Cortex | 3.1 |
| Cuneal Cortex | 4.1 |
| Lingual Gyrus | 9.0 |
| Temporal Occipital Fusiform Cortex | 2.7 |

| Cluster #2 | Average Atlas Probability Within Cluster |
|---|---|
| Insular Cortex | 4.8 |
| Precentral Gyrus | 15.4 |
| Postcentral Gyrus | 16.7 |
| Central Opercular Cortex | 13.0 |

| Cluster #1 | Average Atlas Probability Within Cluster |
|---|---|
| Precentral Gyrus | 20.0 |
| Postcentral Gyrus | 36.8 |

Supplemental Table 10. Atlas query of conjunction of Right Lobule VI - Crus I connectivity results. For large clusters, sub-region labels are listed below. COG: Center of Gravity.

| ClusterIndex | Voxels | Max T-Values | COG X (mm) | COG Y (mm) | COG Z (mm) | COG Label |
|---|---|---|---|---|---|---|
| 16 | 12360 | 50 | 10.6 | -59.4 | -18.5 | 88% Right V |
| 15 | 11542 | 15.4 | -44.5 | -19.7 | 35.8 | 42% Postcentral Gyrus |
| 14 | 1805 | 9 | 59.4 | -10.3 | 21.7 | 38% Postcentral Gyrus |
| 13 | 1393 | 10.4 | -1.56 | 55.9 | -3.89 | 44% Frontal Pole |
| 12 | 699 | 7.01 | -38.6 | -74.9 | 31.5 | 42% Lateral Occipital Cortex, superior division |
| 11 | 648 | 9.8 | -4.75 | -7.44 | 51 | 29% Supplemental Motor Area |
| 10 | 630 | 8.18 | -43.2 | 33.4 | 17.7 | 64% Area 9/46V |
| 9 | 432 | 6.59 | -51.7 | -63 | 0.58 | 35% Middle Temporal Gyrus, temporooccipital part |
| 8 | 407 | 7.4 | -27.7 | -35.4 | -19.1 | 69% Temporal Fusiform Cortex, posteror division |
| 7 | 383 | 6.79 | 47.9 | -63.2 | 27.2 | 66% Lateral Occipital Cortex, superior division |
| 6 | 117 | 7.06 | 59.1 | -6.54 | -13.3 | 31% Middle Temporal Gyrus, anterior division |
| 5 | 80 | 7.08 | -21.3 | -21.1 | -15.2 | 76% Left Hippocampus |
| 4 | 61 | 6 | 28.8 | 2.55 | -14 | 44% Right Amygdala |
| 3 | 53 | 7.37 | -15.6 | -19.2 | 4.48 | 97% Left Thalamus |
| 2 | 47 | 5.96 | -34.3 | -9.63 | -31.8 | 25% Temporal Fusiform Cortex |
| 1 | 38 | 5.49 | 25.4 | -21.8 | -15.3 | 85% Right Hippocampus |

-----------------------------------------

Structures to which each cluster belongs to:

| Cluster #16 | Average Atlas Probability Within Cluster |
|---|---|
| Cingulate Gyrus, posterior division | 4.1 |
| Precuneous Cortex | 12.9 |
| Right V | 4.9 |
| Right VI | 8.5 |

| Cluster #15 | Average Atlas Probability Within Cluster |
|---|---|
| Precentral Gyrus | 11.2 |
| Postcentral Gyrus | 14.6 |

| Cluster #14 | Average Atlas Probability Within Cluster |
|---|---|
| Area 6v | 10.5 |
| Precentral Gyrus | 14.0 |
| Postcentral Gyrus | 18.5 |
| Central Opercular Cortex | 10.7 |

| Cluster #13 | Average Atlas Probability Within Cluster |
|---|---|
| FPm (Frontal Pole medial - ventral) | 16.2 |
| Area 10 (dorsal) | 21.7 |
| Frontal Medial Cortex | 18.4 |
| Paracingulate Gyrus | 14.7 |

| Cluster #12 | Average Atlas Probability Within Cluster |
|---|---|
| Lateral Occipital Cortex, superior division | 63.6 |

| Cluster #11 | Average Atlas Probability Within Cluster |
|---|---|
| Precentral Gyrus | 12.4 |
| Juxtapositional Lobule Cortex | 39.2 |
| Cingulate Gyrus, anterior division | 11.8 |

| Cluster #10 | Average Atlas Probability Within Cluster |
|---|---|
| Area 9/46V | 31.9 |
| Area 44d | 20.0 |
| IFS (Inferior Frontal Sulcus) | 37.3 |
| Area 46 | 13.0 |

| Cluster #9 | Average Atlas Probability Within Cluster |
|---|---|
| Middle Temporal Gyrus, temporooccipital part | 16.6 |
| Inferior Temporal Gyrus, temporooccipital part | 8.8 |
| Lateral Occipital Cortex, inferior division | 38.8 |

| Cluster #8 | Average Atlas Probability Within Cluster |
|---|---|
| Left V | 11.1 |
| Parahippocampal Gyrus, posterior division | 23.3 |
| Temporal Fusiform Cortex, posterior division | 31.5 |
| Temporal Occipital Fusiform Cortex | 6.2 |

| Cluster #7 | Average Atlas Probability Within Cluster |
|---|---|
| Angular Gyrus | 11.7 |
| Lateral Occipital Cortex, superior division | 49.1 |

| Cluster #6 | Average Atlas Probability Within Cluster |
|---|---|
| Superior Temporal Gyrus, anterior division | 16.1 |
| Superior Temporal Gyrus, posterior division | 18.3 |
| Middle Temporal Gyrus, anterior division | 22.2 |
| Middle Temporal Gyrus, posterior division | 18.6 |

| Cluster #4 | Average Atlas Probability Within Cluster |
|---|---|
| Insular Cortex | 11.9 |
| Right Amygdala | 17.7 |

| Cluster #2 | Average Atlas Probability Within Cluster |
|---|---|
| Parahippocampal Gyrus, anterior division | 18.4 |
| Temporal Fusiform Cortex, anterior division | 29.0 |
| Temporal Fusiform Cortex, posterior division | 21.5 |

Supplemental Table 11. Atlas query of conjunction of Left Lobule VI - Crus II connectivity results. For large clusters, sub-region labels are listed below. COG: Center of Gravity.

| ClusterIndex | Voxels | Max T-Value | COG X (mm) | COG Y (mm) | COG Z (mm) | COG Label |
|---|---|---|---|---|---|---|
| 5 | 34521 | 49.8 | 8.46 | -39.7 | 8.39 | 15% Cingulate Gyrus, posterior division |
| 4 | 1502 | 13.1 | 36.3 | 42.8 | 25.6 | 100% Area 46 |
| 3 | 818 | 9.33 | -47.4 | 7.49 | 2.25 | 22% Central Opercular Cortex |
| 2 | 705 | 9.32 | -32.5 | 44.5 | 29.2 | 61% Area 9/46D |
| 1 | 52 | 6.49 | 14.5 | -30.8 | 42.3 | 34% Precentral Gyrus |

------------------------------------------

Structures to which each cluster belongs to:

| Cluster #5 | Average Atlas Probability Within Cluster |
|---|---|
| Precentral Gyrus | 5.2 |
| Lateral Occipital Cortex, superior division | 5.0 |
| Lateral Occipital Cortex, inferior division | 4.2 |

| Cluster #4 | Average Atlas Probability Within Cluster |
|---|---|
| Cluster5 (Area 9/46D) | 19.8 |
| Cluster6 (Area 9/46V) | 15.9 |
| Area 46 | 23.0 |

| Cluster #3 | Average Atlas Probability Within Cluster |
|---|---|
| Insular Cortex | 14.3 |
| Precentral Gyrus | 11.1 |
| Frontal Operculum Cortex | 9.5 |
| Central Opercular Cortex | 14.1 |

| Cluster #2 | Average Atlas Probability Within Cluster |
|---|---|
| Cluster5 (Area 9/46D) | 36.0 |
| Cluster7 (Area 46) | 18.6 |

| Cluster #1 | Average Atlas Probability Within Cluster |
|---|---|
| Precentral Gyrus | 18.9 |
| Postcentral Gyrus | 6.6 |
| Cingulate Gyrus, posterior division | 26.8 |
| Precuneous Cortex | 8.0 |

Supplemental Table 12. Atlas query of conjunction of Right Lobule VI - Crus I connectivity results. For large clusters, sub-region labels are listed below. COG: Center of Gravity.

| ClusterIndex | Voxels | MAX | COG X (mm) | COG Y (mm) | COG Z (mm) | COG Label |
|---|---|---|---|---|---|---|
| 11 | 14245 | 14.8 | -33.3 | -11 | 44.7 | 21% Precentral Gyrus |
| 10 | 6792 | 49 | 22.9 | -58 | -25.7 | 89% Right VI |
| 9 | 1522 | 13.3 | -32.2 | 44.7 | 26.4 | 56% Area 9/46D |
| 8 | 342 | 6.54 | 34 | 47.7 | 26.9 | 100% Area 46 |
| 7 | 292 | 8.87 | -35.7 | -63.2 | -25.9 | 86% Left Crus I |
| 6 | 161 | 6.61 | -56.6 | -41.5 | 25.2 | 31% Supramarginal Gyrus, posterior division |
| 5 | 136 | 5.87 | 9.91 | -91.6 | 17 | 56% Occipital Pole |
| 4 | 103 | 6.47 | 21.4 | 3.07 | 69.5 | 51% Superior Frontal Gyrus |
| 3 | 48 | 6.27 | -35.5 | -55.9 | -51 | 75% Left VIIb |
| 2 | 47 | 6.23 | -9.36 | -56.1 | -31.7 | 10% Left IX |
| 1 | 37 | 6 | 61 | 4.76 | 23.2 | 100% Area 6v |

------------------------------------------

Structures to which each cluster belongs to:

| Cluster #11 | Average Atlas Probability Within Cluster |
|---|---|
| Precentral Gyrus | 11.5 |
| Postcentral Gyrus | 10.6 |

| Cluster #10 | Average Atlas Probability Within Cluster |
|---|---|
| Lingual Gyrus | 5.6 |
| Temporal Occipital Fusiform Cortex | 4.3 |
| Occipital Fusiform Gyrus | 4.4 |

| Cluster #9 | Average Atlas Probability Within Cluster |
|---|---|
| Area 9/46D | 27.1 |
| Area 9/46V | 12.4 |
| Area 46 | 25.5 |

| Cluster #8 | Average Atlas Probability Within Cluster |
|---|---|
| Area 9/46D | 26.3 |
| Area 46 | 27.6 |

| Cluster #6 | Average Atlas Probability Within Cluster |
|---|---|
| Supramarginal Gyrus, anterior division | 8.0 |
| Supramarginal Gyrus, posterior division | 26.0 |
| Parietal Operculum Cortex | 18.4 |
| Planum Temporale | 9.7 |

| Cluster #5 | Average Atlas Probability Within Cluster |
|---|---|
| Cuneal Cortex | 9.1 |
| Supracalcarine Cortex | 4.7 |
| Occipital Pole | 48.3 |

| Cluster #4 | Average Atlas Probability Within Cluster |
|---|---|
| Supplementary Motor Area | 10.2 |
| Anterior PMd (Premotor dorsal) | 12.0 |

| Cluster #1 | Average Atlas Probability Within Cluster |
|---|---|
| Area 6v | 64.9 |
| Precentral Gyrus | 58.5 |
